# “Multiple human adipocyte subtypes and mechanisms of their development”

**DOI:** 10.1101/537464

**Authors:** So Yun Min, Anand Desai, Zinger Yang, Agastya Sharma, Ryan M.J. Genga, Alper Kucukural, Lawrence Lifshitz, René Maehr, Manuel Garber, Silvia Corvera

## Abstract

Human adipose tissue depots perform numerous diverse physiological functions, and are differentially linked to metabolic disease risk, yet only two major human adipocyte subtypes have been described, white and “brown/brite/beige.” The diversity and lineages of adipocyte classes have been studied in mice using genetic methods that cannot be applied in humans. Here we circumvent this problem by studying the fate of single mesenchymal progenitor cells obtained from human adipose tissue. We report that a minimum of four human adipocyte subtypes can be distinguished by transcriptomic analysis, specialized for functionally distinct processes such as adipokine secretion and thermogenesis. Evidence for the presence of these adipocytes subtypes in adult humans is evidenced by differential expression of key adipokines leptin and adiponectin in isolated mature adipocytes. The human adipocytes most similar to the mouse “brite/beige” adipocytes are enriched in mechanisms that promote iron accumulation and protect from oxidative stress, and are derived from progenitors that express high levels of cytokines such as *IL1B*, *IL8*, *IL11* and the *IL6* family cytokine *LIF*, and low levels of the transcriptional repressors *ID1* and *ID3*. Our finding of this adipocyte repertoire and its developmental mechanisms provides a high-resolution framework to analyze human adipose tissue architecture and its role in systemic metabolism and metabolic disease.

## INTRODUCTION

Adipose tissue plays critical roles in systemic metabolism, through multiple functions that include lipid storage and release, as well as secretion of cytokines such as leptin and adiponectin which control satiety and energy utilization [1]. In addition to its metabolic roles, adipose tissue plays a pivotal role in thermoregulation, both by providing an insulating layer under the skin and by the production of heat [2]. Other functions of adipose tissue include mechanical and immune-protective roles in multiple sites including the mesentery [3].

Currently, three functionally distinct adipocyte subtypes in mammals are recognized, which are best defined in the mouse [4]. White adipocytes are lipogenic and responsible for lipid storage and release; brown adipocytes form a defined depot restricted to the interscapular region and contain high basal levels of the mitochondrial uncoupler UCP1, required for their thermogenic function; and “brite/beige” adipocytes are interspersed within white adipose tissue and express UCP1 in response to stimulation. Lineage tracing and gene expression studies point to distinct developmental origins for these adipocyte subtypes [5, 6]. In adult humans, no specific depot is solely composed of UCP1-containing adipocytes, but UCP-1 positive cells can be found interspersed within supraclavicular, paravertebral, and perivascular adipose tissue [7], resembling the “brite/beige” phenotype. Thus, in humans, only two adipocyte types, UCP-1 positive (thermogenic) and UCP-1 negative (white) have been recognized to exist.

How two cell types can accomplish the numerous functions of human adipose tissue is unclear. Increased functionality of adipose tissue may be conferred by the different connections to vascular and neural networks present in distinct depots [4], which could play a role in diversifying adipocyte functions. For example, significant differences in lipolytic capacity and cytokine secretion profiles are seen between depots that are composed of ostensibly similar adipocyte types [8–10]. Alternatively, additional yet uncharacterized developmentally and epigenetically distinct adipocyte subtypes that are not readily distinguished by their morphology may exist.

Recent advances in the ability to study single-cell transcriptomes can be leveraged to better understand the cellular composition of adipose tissue [11–13]. Single-cell transcriptomics of non-mature adipocytes in mouse fat has revealed specific populations of adipocyte stem cells primed to expand in response to adrenergic stimulation [13], stromal cells capable of inhibiting adipogenesis [11], and cells with enhanced adipogenic or fibro-inflammatory properties [14]. The use of single-cell transcriptomics to define mature adipocyte subtypes is complicated by the large size and high buoyancy of these cells, and the fact that the extensive proteolytic digestion required to isolate them from the adipose tissue compromises their viability and alters their phenotype [15]. While elegant solutions to these limitations have been developed [16], they are dependent on transgenic approaches and are thus less amenable to application for human studies.

As an alternative approach to studying human adipocyte subtypes and their development, we leveraged a recent finding by our laboratory that allows us to generate large numbers of mesenchymal progenitor cells from human adipose tissue with little loss of multipotency [17–19]. Many of these progenitor cells differentiate into large, long-lived unilocular adipocytes [18], and notably, some correspond to thermogenic adipocytes, as they respond to stimulation by inducing UCP1 and other brown adipose tissue markers [19]. To identify the distinguishing features of progenitor cells that give rise to adipocytes, and of the adipocyte subtypes generated from these progenitors, we obtained transcriptomic profiles before and after adipogenic induction, and before and after thermogenic stimulation of clonal cell populations derived from single mesenchymal progenitor cells.

Our results uncovered a minimum of four distinct adipocyte subtypes differing in gene expression profiles associated with distinct, recognizable adipocyte metabolic functions. Progenitor cells that give rise to each adipocyte subtype express distinct complements of transcriptional regulators and cytokines, many of which can explain existing results of the effects of genetic ablation or overexpression of these factors on adipose tissue function. Thus, our results provide a high-resolution perspective on the complexity of human adipose tissue, and propose tangible models that can be further tested to better understand human adipose tissue function and systemic metabolism.

## RESULTS

### Morphological evidence for distinct adipocyte subtypes

To explore the mechanisms by which human adipocyte subtypes originate, we developed an approach summarized in Figure 1a and detailed in the Methods. Cells sprouting from human adipose tissue explants are plated as single cells, and proliferation allowed until near confluence, when they are split 1:3 into three wells of a 384 well multiwell plate. One well is maintained in a non-differentiated state (C), two wells are subjected to adipose differentiation (M), and one of the differentiate wells (F) is stimulated with Forskolin (Fsk) for the last 3 days of culture. Wells are then imaged and RNA extracted. Phase images of clones after differentiation and Fsk activation revealed three main subtypes. One consisted of cells that proliferated to confluence, but fail to accumulate lipid droplets in response to differentiation, and were therefore not considered further in this study (Figure 1b). A second subtype of cells accumulated lipid droplets, but were not visibly affected by Fsk (Figure 1c), suggesting a weak response to the lipolytic effects of this drug. A third subtype of cells accumulated lipid droplets and responded to Fsk with visible changes in droplet size and number (Figure 1d), suggesting a strong lipolytic response. To determine whether these differences were explained by expected stochastic variation, or whether they reflect the existence of true distinct adipocyte subtypes, we analyzed the frequency distributions of droplet size, measured after image processing as exemplified (Figure 1 b-d, lower panels). Aggregated values for lipid droplet size from all clones (Figure 1e), produced a monophasic histogram representative of the normal variation of adipocyte droplet sizes in these cells collectively. However, when we plotted the mean droplet size per clone, we obtained a non-monophasic distribution (Figure 1f), revealing the existence of adipocyte subtypes that differ in mean droplet size. We then quantified the responsiveness of each clone by comparing the size of lipid droplets before and after Fsk treatment (Figure 1g). Again, Fsk responsiveness was not represented by a monophasic distribution, as the histogram of the values (droplet size in MDI+Fsk as % of MDI only value) contained at least two peaks, a major peak centered around 100% (no response) and another at around 60% (decrease of lipid droplet size to 60% of non-stimulated value) (Figure 1h). These results suggest that at least two adipocyte subtypes exist, based on their mean droplet size and differential responsiveness to Fsk.

**Figure 1.**
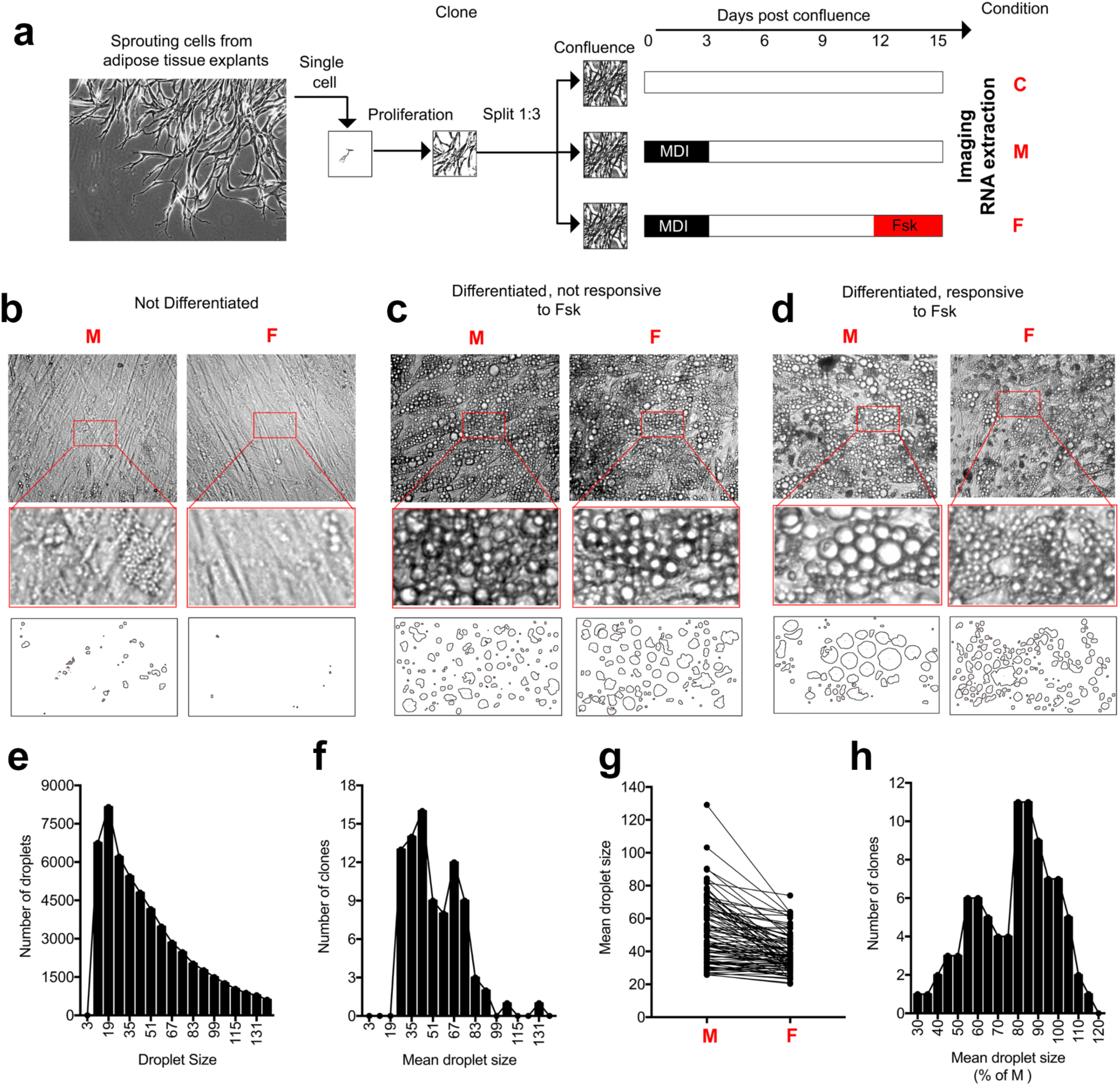
Single mesenchymal progenitors give rise to morphologically different adipocyte subtypes. **a.** Mesenchymal progenitor cells from adipose tissue are induced to proliferate under pro-angiogenic conditions in a 3-dimensional hydrogel. Single cells are isolated, and proliferation allowed until near confluence, when they are split 1:3 into three wells of a 384 well multiwell plate. One well is maintained in a non-differentiated state (C), two wells are subjected to adipose differentiation (M), and one of the differentiate wells is stimulated with Forskolin (Fsk) for the last 3 days of culture (F). Wells are then imaged and RNA extracted. **b.** Example of a clone that failed to undergo adipose differentiation; **c**. Example of a clone that underwent adipocyte differentiation but was unresponsive to Fsk as assessed by lipid droplet size; **d**. Example of a clone that underwent adipocyte differentiation and responded to Fsk with decrease in lipid droplet size. **e**. Frequency distribution of all lipid droplets measured in all clones that underwent adipocyte differentiation. **f**. Frequency distribution of the mean droplet size per clone in clones that underwent adipocyte differentiation. **g**. Measurement of mean droplet sizes per clone comparing M and F conditions. **h**. Frequency distribution of responsiveness to Fsk, where mean lipid droplet size is expressed as percent of droplet size in M for each specific clone.

### Identification of transcriptionally distinct adipocyte subtypes

We then asked whether the morphological evidence for distinct human adipocyte subtypes is supported by differences in gene expression. We extracted RNA from 156 samples comprising non-differentiated (C), differentiated (M) and differentiated plus Fsk-stimulated (F) conditions for each of the 52 clones obtained (Figure 1a). Principal component analysis revealed clear segregation of C, M and F conditions (Figure 2a), verifying that differentiation and Fsk stimulation had major effects on gene expression in these clones. Interestingly, a broader spread is seen in the first principal component of M and F compared to C conditions, indicating generation of divergent transcriptional landscapes following induction of differentiation and thermogenic stimulation.

**Figure 2.**
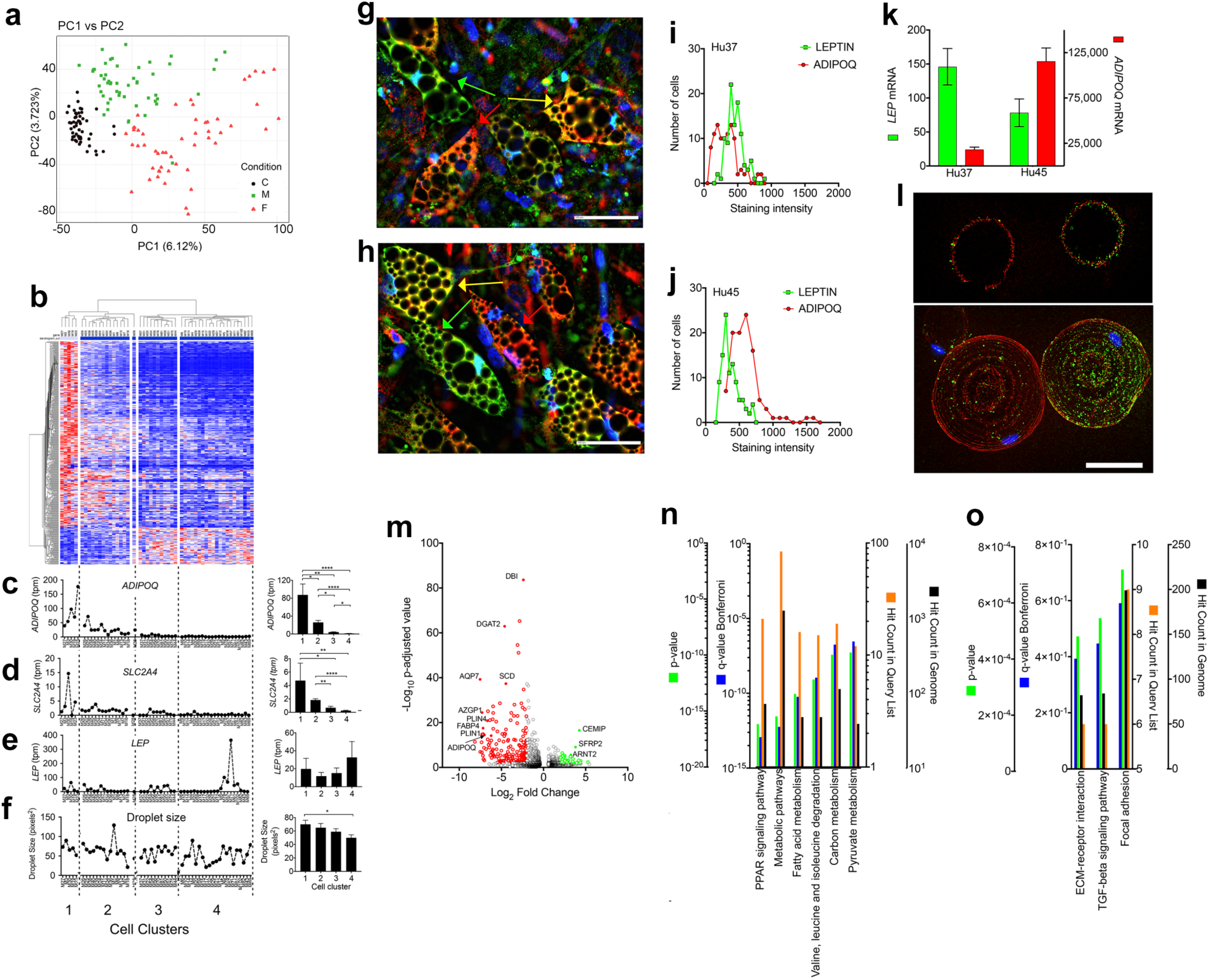
Adipocyte progenitors give rise to distinct adipocyte subtypes with different functional properties. **a.** Principal component analysis of 52 clones analyzed under non-differentiated (C, black symbols), differentiated (M, green symbols) and differentiated plus (Fsk) stimulated (F, red symbols) conditions. **b.** Unsupervised hierarchical clustering of TPM values for 447 genes (Supplementary Table 1) correlated with lipid droplet size. Each row is independently scaled from blue (lower) to red (higher) values. **c-e.** TPM values for the genes indicated in the y axis for the clones in the x-axis Panels on the left are the mean and SEM of TPM values from each cluster defined by the heat map above. **f.** Mean droplet size for the clones in the x-axis. Panels on the left are the mean and SEM for each cluster as defined by the heat map above. Statistical comparison between all columns was done using the Kruskal-Wallis test corrected for multiple comparisons by controlling the false discovery rate using the Benjamini, Krieger and Yekutieli test as implemented in GraphPad Prism 7. *=p<0.05; **=p<0.01; ****=p<0.0001. **g,h.** Immunofluorescence staining of differentiated adipocytes from two individuals using antibodies to leptin and adiponectin, and Hoechst for nuclear staining. Arrows indicate adipocytes expressing mostly leptin (green), mostly adiponectin (red) or both (yellow). **i,j.** Frequency distribution of staining intensities per cell for leptin (green) or adiponectin (red) from 100 cells in fields exemplified in g and h. **k.** qRT-PCR of *LEP* and *ADIPOQ* in cultures exemplified in g and h. Values are expressed as fold difference relative to the lowest detectable value. **l.** Immunofluorescence staining of primary human adipocytes isolated by collagenase digestion. Green is leptin and red is adiponectin. Top panel is a single optical plane and bottom is the projection of all optical planes comprising the 3-dimensional volume of the cells. **m.** Volcano plot comparing cluster 1 and cluster 4. Red and green symbols are genes enriched in cluster 1 or cluster 4, respectively. **n,o.** KEGG pathway enrichment of enriched genes in cluster 1 or cluster 4, respectively.

We then analyzed the transcriptomes of clones in the M condition using hierarchical clustering. As a first approach, we sought to focus on genes that were associated with key properties of adipocytes. Thus, we clustered genes that correlated with lipid droplet size (as assessed in Figure 1) with a Pearson rank order correlation coefficient higher than 0.5 or lower than −0.5 (Supplementary Table 1). Unsupervised hierarchical clustering of these 447 genes resulted in 4 major clusters (Figure 2b) indicating 4 distinct adipocyte subtypes. To determine how these adipocyte subtypes related to classical adipose tissue traits, we looked at the expression levels of the canonical adipocyte genes *ADIPOQ* (Adiponectin), *SLC2A4* (GLUT4) and *LEP* (Leptin) within these clusters. Clones in Clusters 1 and 2 expressed the highest levels of *ADIPOQ* and *SLC2A4* (Figure 2 c,d), but surprisingly the highest expression of *LEP* was within cluster 4, which displayed lower expression levels of *ADIPOQ* and GLUT4 (Figure 2e). Mean lipid droplet size in Cluster 4 was also significantly smaller compared to Cluster 1 (Figure 2f). These results reveal the existence of adipocyte subtypes with distinct transcriptomic states associated with key adipocyte functions.

**Table 1:**
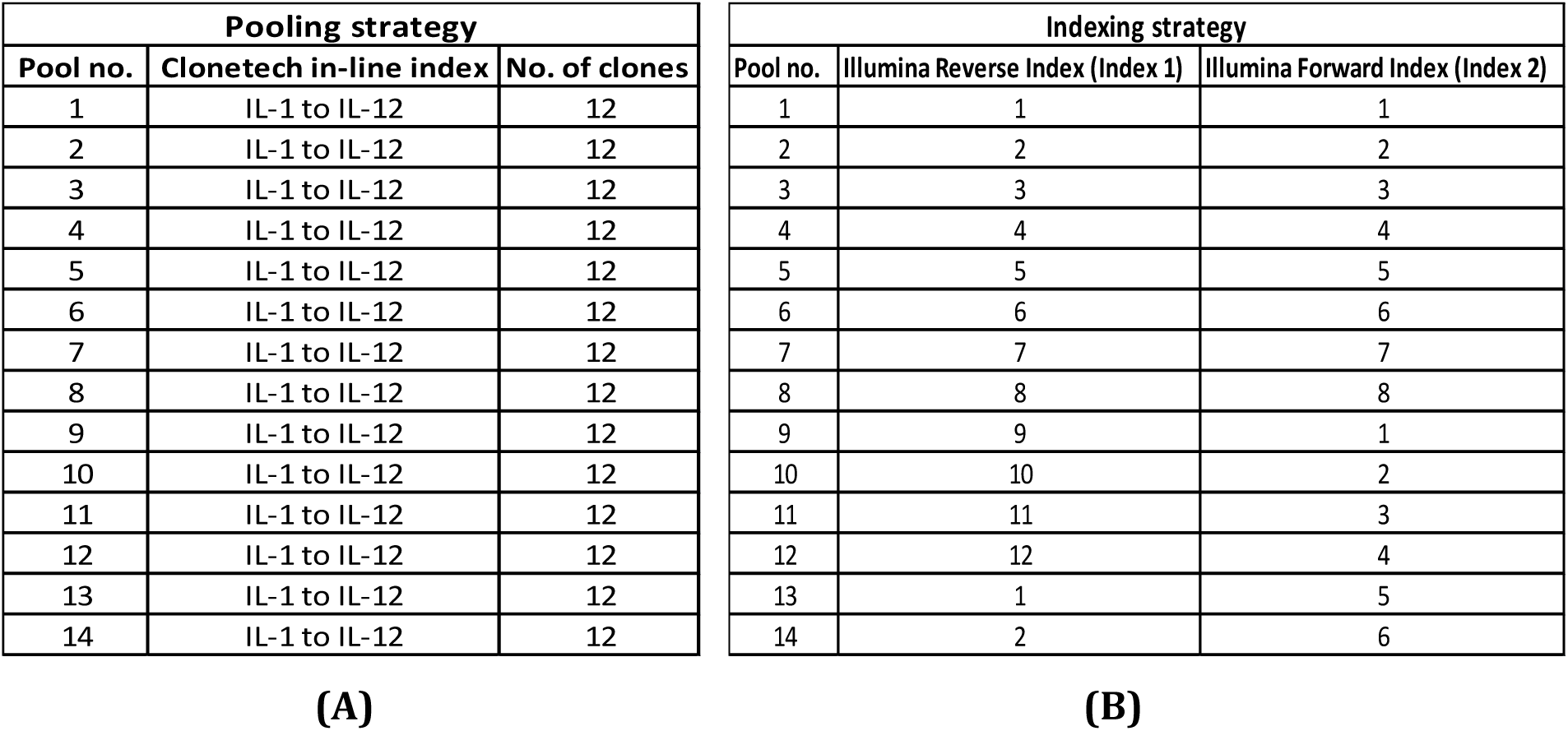
Pooling and indexing strategy for 168 samples grouped in fourteen pools.

To verify the robustness of these results, we performed an additional, unsupervised clustering analysis of the M samples using the top 1500 genes displaying the highest variance. This analysis produced 5 distinct clusters, overlapping with the clusters generated by hierarchical clustering of genes correlating with lipid droplet size (Supplementary Figure 2). Importantly, *LEP* and *ADIPOQ*-expressing adipocytes were again segregated in two different clusters.

To test whether the differences in *ADIPOQ* and *LEP* gene expression between adipocyte subtypes would be reflected by the levels of expression of their encoded proteins, we performed immunofluorescence on adipocytes differentiated from unsorted progenitors obtained from two independent subjects. In both cases, cells expressing LEP and very little ADIPOQ, and vice versa, could be readily found (Figure 2g,h). Quantitative analysis of expression was done on 100 cells from each subject. Histograms of these values revealed very different frequency distributions of cells expressing ADIPOQ or LEP (Figure 2, i,j), consistent with these proteins being expressed by different adipocyte types. In addition, differences in the frequency distributions between the two subjects were observed (Figure 2, compare panel i with panel j). Notably, the inter-subject difference in distribution, i.e. the relative number of cells expressing each protein, correlated with the difference in overall expression of *ADIPOQ* and *LEP* mRNA between the two subjects (Figure 2k).

To explore whether the development of adipocytes preferentially expressing *ADIPOQ* or *LEP* mRNA occurs in vivo, we stained for LEP and ADIPOQ in isolated primary adipocytes obtained by collagenase digestion of human subcutaneous adipose tissue. We found that expression in mature unilocular adipocytes is heterogenous (Figure 2l) and that some adipocytes expressed high levels of only one of these proteins. Interestingly, staining of adipocytes from mouse epididymal adipose tissue has already been remarked as being similarly heterogeneous, with some cells expressing no detectable leptin [20]. While immunostaining alone cannot distinguish between stochastic variations in protein levels and the existence of true cell subtypes, the consistency between gene expression levels in clones and single-cell immunostaining of primary cultured and freshly isolated cells support the existence of distinct subtypes of white adipocytes specialized for production of specific adipokines.

To enumerate the genes that define these specific clusters, and to delineate pathways enriched by these genes, we performed differential expression and pathway enrichment analysis between Clusters 1 and 4 as defined in Figure 2b. We find 1308 differentially expressed genes with a p-adjusted value < 0.05. Of these, 174 genes were more highly expressed by > 5-fold in Cluster 1, and 48 genes were more highly expressed > 5-fold in Cluster 4 (Figure 2m and Supplementary Table 2). Pathway enrichment analysis of genes enriched in Cluster 1 included PPARγ signaling and lipogenesis pathways, containing genes such as *DGAT2*, *FABP4*, *PLIN1* and *PLIN4* (Figure 2n and Supplementary table 3), and genes enriched in Cluster 4 were associated with extracellular matrix interaction and TGFβ signaling (Figure 2o and Supplementary table 3). Individual genes that were significantly enriched in Cluster 4 included *SFRP2*, an important regulator the Wnt signaling pathway which is central to adipogenesis [21].

### Identification and characterization of the “brite/beige” adipocyte subtype

We then asked whether any of the clusters would correspond to adipocytes of the “brite/beige” phenotype, as defined by the ability to respond to Fsk stimulation with expression of canonical thermogenic genes. Adipocytes differentiated from heterogeneous progenitor cultures were highly responsive to Fsk stimulation, with strong induction of *UCP1, DIO2 and CIDEA* after acute (6-24h) and chronic (24h to 7 days) exposure (Figure 3a), and with high expression of UCP1 protein after chronic exposure (Figure 3b). Notably, immunofluorescence staining of UCP1 was heterogeneous, consistent with “brite/beige” adipocytes representing a subtype within the population (Figure 3b). We then analyzed the effect of Fsk on lipid droplet size and expression of thermogenic genes in each clone. Most clones responded to Fsk with a decrease in adipocyte droplet size (Figure 3c), as expected from stimulation of lipolysis in response to the drug. In addition, Fsk stimulated *ADIPOQ* expression in Clusters 1 and 2, and inhibited *LEP* expression in Cluster 4 (Figure 3d,e), which are responses previously seen in this cell population [19]. The brown adipose tissue genes *DIO2* and *CIDEC* were more strongly induced in Cluster 2 (Figure 3f,g), suggesting this cluster as corresponding to the “brite/beige” adipocyte subtype.

**Figure 3.**
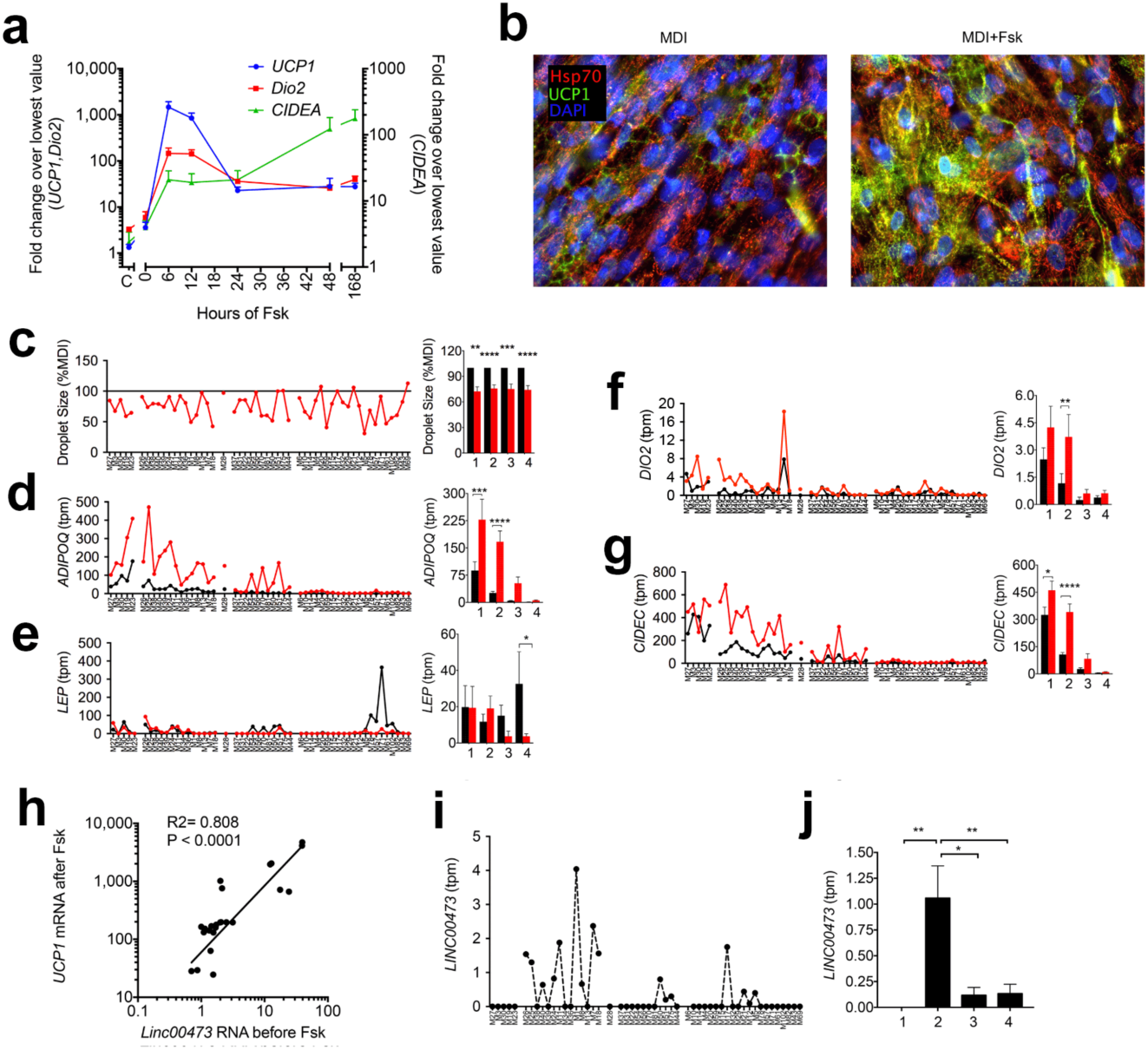
Cluster 2 corresponds to “brite/beige” adipocytes. **a**. RT-PCR of thermogenic genes in non-differentiated (C) and differentiated cells at different times of exposure to Fsk. Values are expressed as fold relative to the lowest value. Ct values for UCP1 were 29-31 at t=0 and decreased to 20-22 at t=6h in response to Fsk. Symbols are means and bars SEM of two independent experiments using cells from two subjects assayed in duplicate. b. Immuofluoresence staining using antibodies to mitochondrial Hsp70 (red) and UCP1 (green) of differentiate cells without or with exposure to Fsk for 7 days. **c-g**. TPM values for the genes indicated in the y-axis for the clones in the x-axis, segregated into clusters as defined in Figure 2. Black symbols and traces are the values for M and red symbols and traces are values for F. Panels on the left are the mean and SEM of TPM values from each cluster. Statistical comparison between M and F in each cluster was done using ANOVA, with correction for multiple comparisons using the Holm-Sidak test as implemented in GraphPad Prism 7. *=p<0.05; **=p<0.01; ***=p<0.001; ****=p<0.0001. **h.** Levels of Linc00473 and UCP1 analyzed by RT-PCR in differentiated adipocytes with and without Fsk stimulation for 6h, and values expressed as fold relative to lowest value. Levels of Linc00473 in non-stimulated cells are plotted against levels of UCP1 after stimulation. Linear regression analysis was performed using GraphPad Prism 7. **i,j**. TPM values for Linc00473 for the clones in the x-axis, segregated into clusters as defined in Figure 2. Bars are the mean and SEM of TPM values from each cluster.

For unknown reasons, *UCP1* reads were detected only in a few clones, despite strong RT-PCR signals (Figure 3a) and accumulation of UCP1 protein (Figure 3a,b). To further test whether Cluster 2 represents “brite/beige” adipocytes, we leveraged recent findings that the levels of the long non-coding RNA, *LINC00473*, highly correlated with *UCP1* levels in “brite/beige” human adipose tissue and in adipocytes derived from this depot [22]. Indeed, we verified this correlation by analyzing the levels of *LINC00473* prior to stimulation by Fsk to the levels of *UCP1* following stimulation in adipocytes differentiated from non-cloned cells from 3 independent subjects (Figure 3h). *LINC00473* was highly enriched in Cluster 2 (Figure 3 i,j), consistent with this cluster representing the “brite/beige” adipocyte subtype.

To define features that distinguish “brite/beige” adipocytes from other subtypes, we performed differential expression analysis between Cluster 2 and the combined Clusters 1, 3 and 4 (Figure 4a). We found 85 genes to be differentially expressed with a p-adjusted value < 0.05. Of these, 12 genes were increased by 1.5-fold or more, and 32 genes were decreased by −1.5-fold or more in Cluster 2 relative to all other clusters (Supplementary Table 4). Pathway enrichment analysis of genes increased in Cluster 2 with a p-adjusted value of 0.1 or lower (Supplementary Table 5) revealed pathways related to carbohydrate, carbon and nitrogen metabolism. It is notable that the gene family of carbonic anhydrases was highly enriched (p-adj = 4.505E-4) by the presence of *CA3* and *CA9* (Figure 4a, Supplementary Table 5). Genes decreased in Cluster 2 corresponded to the ferroptosis and mineral absorption pathways (Figure 4c), which are defined by the gene *SLC40A1* (Ferroportin, *FPN1*) (Figure 4a, Supplementary Table 5), the only mechanism known to mediate iron egress from cells.

**Figure 4.**
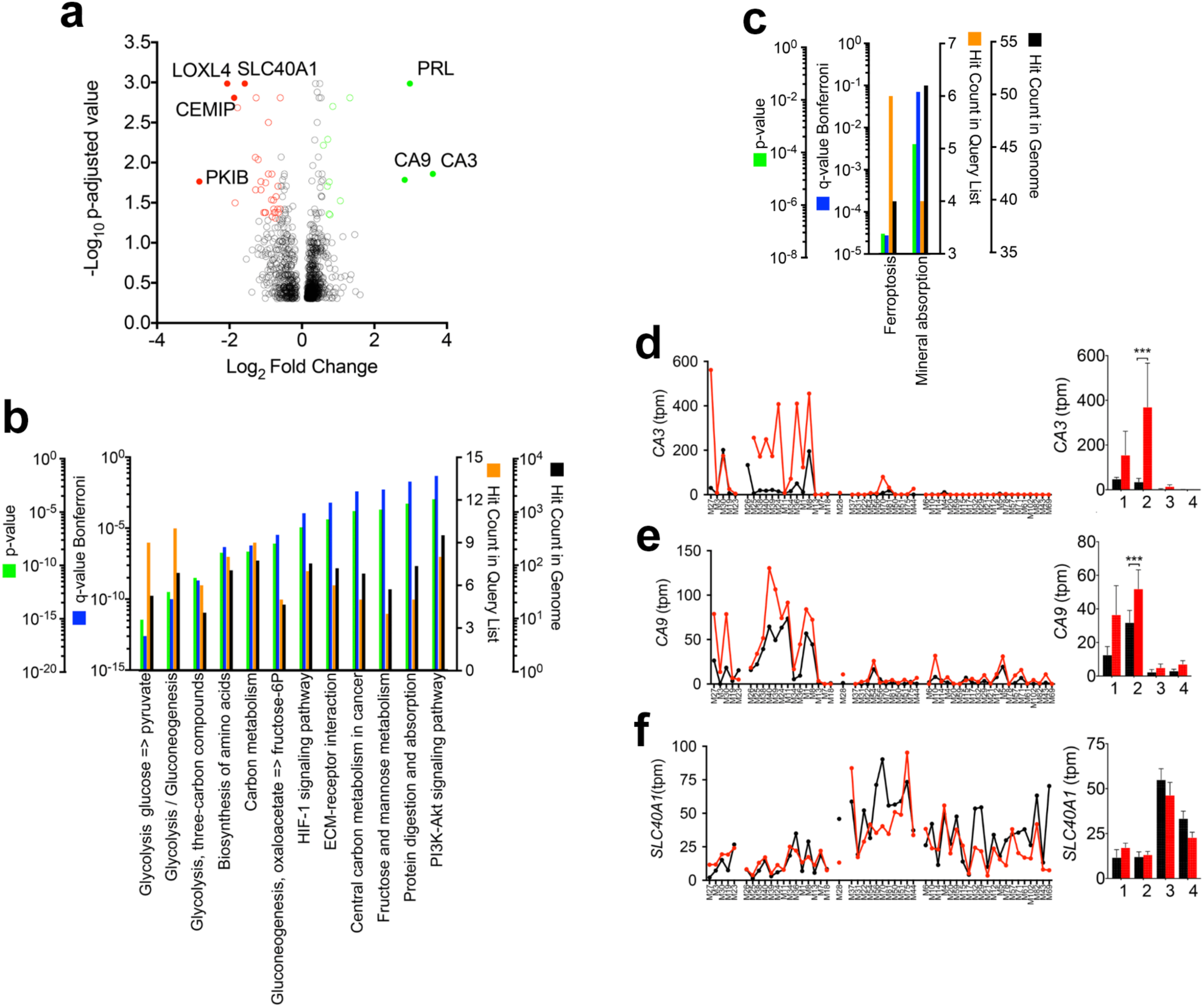
Differential gene expression analysis identifies genes that characterize human “brite/beige” adipocytes. **a.** Volcano plot of differential expression analysis results comparing Cluster 2 with all other clusters. **b,c.** KEGG pathway analysis of genes increased or decreased in Cluster 2 relative to all other clusters. **d**-**f.** TPM values for the genes indicated in the y-axis for the clones in the x-axis, segregated into clusters as defined in Figure 2. Black symbols and traces are the values for M and red symbols and traces are values for F. Panels on the left are the mean and SEM of TPM values from each cluster. Statistical comparison between M and F in each cluster was done using ANOVA, with correction for multiple comparisons using the Holm-Sidak test as implemented in GraphPad Prism 7. ***=p<0.001.

We then examined the effects of Fsk on the levels of *CA3*, *CA9*, and *SLC40A1*/*FPN1* in all clones. Notably, Fsk stimulated expression of *CA3* in Cluster 2, but not in other clusters (Figure 4d). Interestingly, *CA3* is thought to be highly tissue-specific for skeletal muscle, consistent with previously observed similarities between thermogenic adipose tissue and muscle [6]. Expression of *CA9* also was stimulated by Fsk, but to a lesser extent (Figure 4e). Fsk did not significantly affect expression of *SLC40A1*/*FPN1*, which remained under-represented in both Clusters 1 and 2 (Figure 4f). Taken together, these results indicate that Cluster 2 comprises “brite/beige” adipocytes, and these adipocytes are primed to increase metabolism, accumulate iron, and resist ensuing oxidative stress.

### Characteristics of progenitor cells that give rise to adipocyte subtypes

The results of differential expression analysis indicate that adipocyte subtypes have different physiological functions; Cluster 1 represents a highly lipogenic subtype, Cluster 2 is a “brite/beige” thermogenic subtype and Cluster 4 is specialized for leptin secretion. To explore the mechanisms by which distinct adipocyte subtypes might be generated, we analyzed the features of the progenitor cells (C condition) that give rise to cells in each cluster. We first compared the progenitors that gave rise to Cluster 1 with all others to define what genes might predict the development of highly lipogenic adipocytes. 36 genes were differentially expressed by 2-fold with a p-adjusted value of 0.05 (Supplementary Table 6). Only 2 of these genes (*EVI2A* and *CCDC69*) were up-regulated in progenitors giving rise to Cluster 1 (Figure 5a). Pathway enrichment analysis of genes depleted in Cluster 1 progenitor cells revealed the KEGG pathway for IL17 signaling, which includes *CXCL6*, *CXCL5*, *CXCL1* (Figure 5b and Supplementary Table 7).

**Figure 5.**
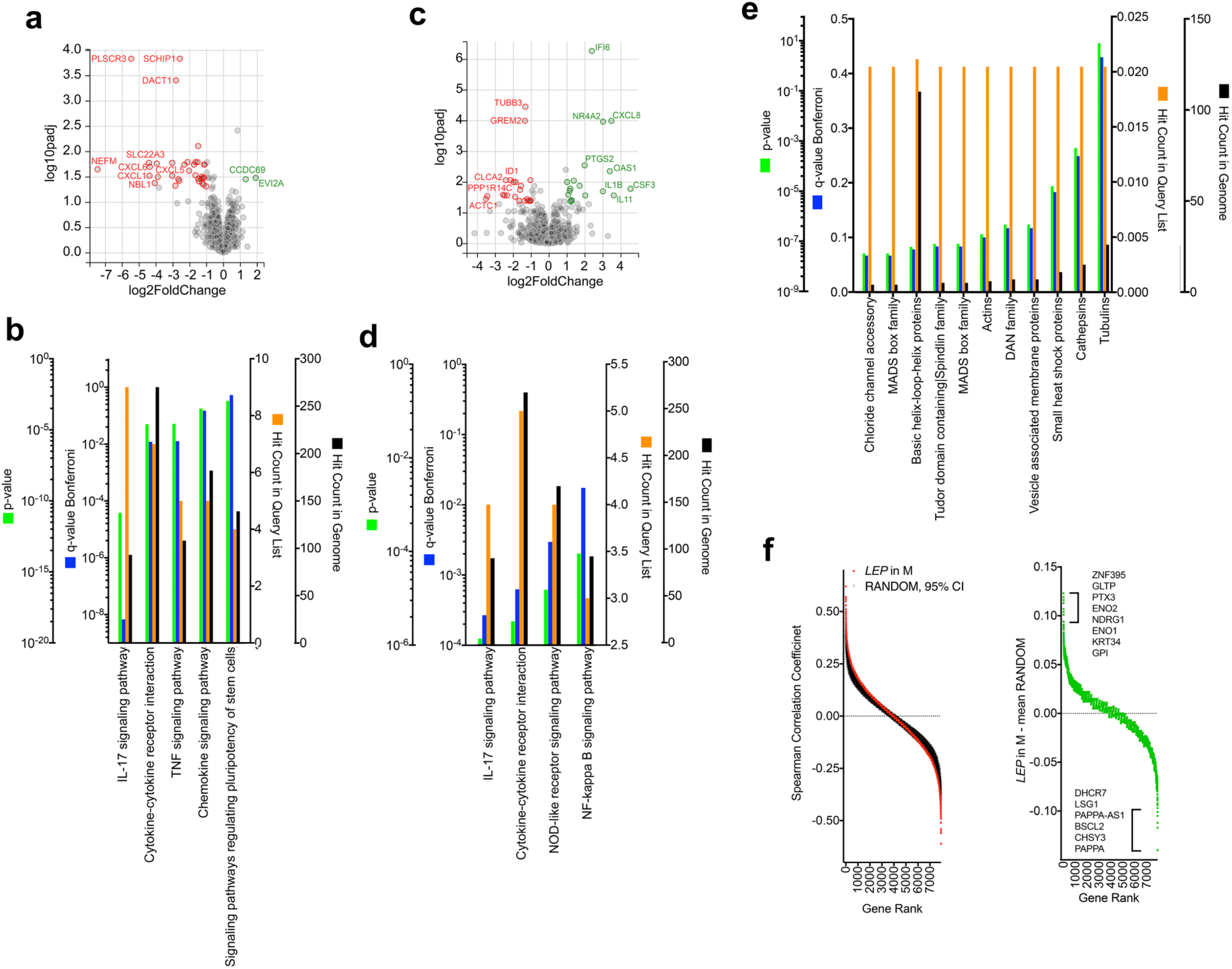
Features of progenitor cells that give rise to adipocyte subtypes. **a.** Volcano plot of differential expression analysis results comparing progenitors corresponding to Cluster 1 with all others. **b.** KEGG pathway analysis of genes under-represented in Cluster 1. **c.** Volcano plot of expression analysis results comparing progenitors corresponding to Cluster 2 with all others. **d.** KEGG pathway analysis of genes over-represented in Cluster 2. **e.** Gene family enrichment of genes under-represented in Cluster 2. **f.** Spearman rank order correlation between genes in C condition and the mean and 95% confidence interval of 10 sets of random numbers (RANDOM) or between genes in C condition and *LEP* levels in M condition. Right panel is the difference between the correlation coefficients seen in the left panel.

To define what genes might predict the development of “brite/beige” adipocytes, we then compared the progenitors that gave rise to Cluster 2 with all others. 39 genes were differentially expressed by 2-fold with a p-adjusted value of 0.05 (Supplementary Table 8). Of these, 18 had higher expression and 21 had lower expression in comparison to all other clones (Figure 5c). Pathway enrichment analysis of genes enriched in Cluster 2 progenitors mapped to the KEGG pathway for IL17 signaling, including *CSF3*, *IL1B*, *PTGS2* and *CXCL8* (Figure 5d, Supplementary Table 9). Other genes significantly enriched in Cluster 2 include *NR4A2*, a cyclic AMP-responsive gene that has been implicated in enhancing UCP1 transcription [23]. The 21 genes under-represented in Cluster 2 were not enriched in any currently specified pathway but mapped significantly to MAD box and basic helix-loop-helix gene families (*MEF2C*, *ID1* and *NPAS1*) (Figure 5e, Supplementary Table 9).

To define what genes might predict the development of adipocytes expressing *LEP*, we compared the progenitors giving rise to Cluster 4 with all others. This comparison did not result in any significantly differentially expressed genes. As an alternative approach, we searched for genes expressed in the C condition that might correlate significantly with *LEP* levels in the M condition. We first defined the spurious level of correlation by calculating the Spearman rank order correlation coefficient with 10 sets of 52 random numbers (to match the number of analyzed clones, generated using the “RAND” function in Excel). The mean and 95% CI of the Spearman correlation coefficient between genes in the C condition and the random number sets was compared with their correlation with *LEP* in M (Figure 5f). The difference between the Spearman correlation coefficient between random numbers and *LEP* values decreased asymptotically (Figure 5f, right panel), and we further analyzed the genes displaying differences larger than 0.05 (Supplementary Tables 10 and 11). The gene most strongly positively correlated was *ZNF395*, which has been previously associated with adipogenesis [24, 25], and pathways most enriched included the glycolysis pathway (genes *ENO1* and *ENO2*) and pathways associated with pathogenic infection which are defined by cytoskeletal genes (Supplementary Table 11). The genes most negatively correlated included PAPPA (pappalysin), which regulates local bioavailability of IGF-1 and is differentially expressed in pre-adipocytes from different depots [26], and BSCL2, which is associated with lipid droplet formation and when mutated gives rise to congenital generalized lipodystrophy. Pathways most enriched in genes negatively correlated with *LEP* expression after differentiation included steroid and terpenoid synthesis pathways (Supplementary Table 11).

## DISCUSSION

The major finding of this study is the existence of multiple distinct human adipocyte subtypes, differing in gene expression profiles related to key adipose tissue functions. These findings expand our understanding of adipocyte heterogeneity from white vs “brown/brite/beige” to subtypes within these categories that can collectively mediate the multiple diverse functions of adipose tissue. Our findings were made by analyzing the trancriptomes of cell populations derived from single human mesenchymal pluripotent cells after subjecting these populations to specific adipogenic and thermogenic stimuli. Among many advantages of this approach, one is that stochastic variations in degree and timing of gene expression displayed by single cell types, such as those seen in populations of 3T3-L1 adipocytes [27–31] are averaged out in the population, and thus adipocyte subtype identity is determined from average population responsiveness to adipogenic and thermogenic stimulation. Our finding of distinct adipocyte phenotypes is consistent with prior observations of heterogeneity within adipocytes from human and mouse adipose tissue [32]. While these prior observations could be potentially attributed to stochastic variation, our results showing underlying gene expression architecture suggest that they reflect the existence of human adipocyte subtypes. Moreover, our findings trace these differences to progenitor cells that give rise to each subtype.

While we expected to find two adipocyte subtypes, corresponding to white and “brite/beige” adipocytes, we were surprised to find additional subtypes differing in key adipocyte functions, such as lipogenic capacity and LEP and ADIPOQ expression (Figure 2e). This finding may help understand the basis for the known differences in LEP expression associated with different depots [33], and the association between systemic LEP levels and degree of adiposity, which correlates but is not fully explained by adipocyte size [34], or lipid content. For example, *LEP* expression is not increased under conditions where adipocyte lipid accumulation is increased due to inhibition of lipolysis [35]. Indeed, mean lipid droplet size in adipocytes preferentially expressing *LEP* (Cluster 4) was significantly smaller compared to other clusters (Figure 2f). We also find that adipocytes specialized for *LEP* expression are highly responsive to Fsk (Figure 3e), suggesting that systemic LEP levels might be defined by the relative number and responsiveness of these cells.

Indeed, levels of *LEP* expression in mixed adipocyte populations is explained by the number of adipocytes expressing high LEP levels, rather than the mean levels of expression per cell (Figure 2i-k). Interestingly, *SFRP2*, a potent antagonist of Wnt signaling, is amongst the most highly differentially expressed genes in these *LEP*-expressing adipocytes. Wnt suppresses adipocyte differentiation, and this effect is antagonized by SFRPs [36]. Moreover, levels of circulating SFRP2 are positively correlated with BMI in humans [37]. Together, these results suggest that adipocytes that express *LEP* can stimulate adipogenesis through regulation of Wnt signaling, and that this mechanism could contribute to the positive relationship between systemic LEP levels and increased adiposity.

Our results also distinguished an adipocyte subtype corresponding to the “brite/beige” phenotype, which is defined by increased mitochondrial biogenesis and uncoupling upon cAMP elevation. We find that “brite/beige” human adipocytes have features that enable them to accommodate increased mitochondrial biogenesis and oxidative stress following stimulation. Most salient are the decrease in *SLC40A1*/*FPN1*, the only known mechanisms for iron efflux [38], and the increased expression of carbonic anhydrases *CA2* and *CA3*, which potently protect cells from oxidative stress [39–41]. Interestingly, CA3 is expressed at much higher levels in human skeletal muscle [41, 42] consistent with the notion that thermogenic adipose tissue and muscle share developmental features [6]. Iron metabolism is tightly associated with human adipose tissue function, as seen by increased levels of genes leading to iron efflux in response to overfeeding and their reversal with weight loss [43, 44]. Iron is a rate limiting and regulatory factor in adipose tissue browning and mitochondrial respiration [45, 46], and thereby is tightly associated with redox stress. The parallel down-regulation of *SLC40A1/FPN1* to increase cellular iron, and up-regulation of *CA2* and *CA3* to protect from redox stress, provide a permissive environment for rapid mitochondrial biogenesis and enhanced respiratory flux. These are further enhanced in response to cAMP as evidenced by the elevation of *CA3* expression in “brite/beige” adipocytes in response to Fsk (Figure 4d).

Our experimental paradigm allows us to explore the differential gene expression landscape of progenitors that give rise to adipocyte subtypes. “Brite/beige” adipocyte progenitors display high expression levels of cytokine genes, which could modulate the local concentrations of immune and stromal cells, which in turn could affect fate determination. Indeed, multiple reports exist of paracrine interactions with immune cells that enhance thermogenic activation [47–49]. In addition to cytokines, progenitors that give rise to “brite/beige” adipocytes display enhanced expression of *PTGS2*, the rate-limiting step in prostaglandin synthesis. *PTGS2* has been directly associated with formation of brown adipocytes in mice [50], and partial *PTGS2* deficiency leads to obesity [51]. The results shown here suggest that the role of this enzyme may be conserved in humans, and that it may contribute to browning through effects on “brite/beige” adipocyte progenitors.

In addition to cytokines, genes that regulate gene expression are differentially expressed in progenitors for distinct adipocyte subtypes. For example, progenitors of Cluster 1, which represents adipocytes with high lipogenic capacity, are highly enriched in *CCDC69*, a coiled-coil domain protein implicated in central spindle assembly [52], which is enriched in human adipose tissue (www.proteinatlas.org/ENSG00000198624-CCDC69/tissue). The 21 genes decreased in Cluster 2 relative to other clones mapped significantly to MAD box and basic helix-loop-helix gene families (*MEF2C*, *ID1* and *NPAS1*). *Id1* depletion results in increased generation of thermogenic adipose tissue in mice, consistent with its lower levels in human progenitors fated for the “brite/beige” phenotype [53, 54].

In summary, our results provide evidence for multiple human adipocyte subtypes specialized for key functions associated with adipose tissue, arising from progenitors that can be distinguished by their complement of expressed cytokines and transcriptional regulatory factors. Many of our results are consistent with published observations of the role of these regulatory factors, and many additional results provide the basis for exploration of additional currently unknown mechanisms of adipocyte differentiation and adipose tissue function. The concept that adipose tissue function results from the ensemble of multiple subtypes of cells, rather than from modulation of function of a single cell type will help deconvolve the complex relationships between adipose tissue biology and metabolic disease.

## ACKNOWLEDGEMENTS

This work was supported by grant DK089101-04 to SC. We acknowledge the use of services from the UMASS Bioinformatics Core, supported by NIH CTSA grant UL1 TR000161-05.

## AUTHOR CONTRIBUTIONS

SC supervised this work. SYM, AD, ZY, AS, RMJG, AK, LL, RM, MG: hypothesis generation, conceptual design, data analysis, and manuscript preparation. SYM, AD, ZY, AS, LL conducted experiments and data analysis.

## DECLARATION OF INTERESTS

The authors declare no competing interests.

## MATERIALS AND METHODS

### Clonal isolation and expansion of cells

Cells were obtained as outlined in a previously published method [19]. Briefly, explants from human subcutaneous adipose tissue were embedded in Matrigel (BD Biosciences Cat. No. 356231) and cultured in EBM-2 medium supplemented with endothelial growth factors (EGM-2MV) (Lonza) for 14 days. Sprouting cells were collected using dispase, plated on 15-cm tissue culture plates and grown to confluency. Media from these plates (25 mL per plate) was collected, spun at 2000 x g for 10 min, filtered through 0.2 μM filter and mixed with FBS (65% Media and 35% FBS). This FBS enriched medium was then mixed with fresh EGM-2MV media in a 1:1 ratio and used for collecting and maintaining sorted single cells.

For single cell sorting, a vial of frozen cells was revived and cells sorted as singlets into 384-well plates using a BSL3 BD FACSAria Cell Sorter (BD Biosciences). Clones that grew to confluency were further split 1:3. Briefly, media was removed, and cells were washed with 1X PBS (100 μL). Trypsin (1X) + 0.05% collagenase (50 μL total) was added and incubated at 37ᵒC until the cells were detached. Conditioned media (45 μL) was added and suspensions divided into 3 wells.

Additional 50 μL of conditioned media was added to each well and cells were grown to confluence. Cells in control group (C condition) were maintained in DMEM + 10% FBS. Adipogenic differentiation of two of the three wells was induced by providing confluent cells with DMEM + 10% FBS supplemented with 0.5 mM 3-isobutyl-1-methylxanthine, 1 μM dexamethasone and 1 μg/ml insulin (MDI). Half of MDI media was removed daily and replaced with media with 2x MDI. After 72h the MDI media was replaced with DMEM + 10% FBS. 50% of the media was replaced with fresh media every 48h until day 14. At that point, one of the two wells was stimulated by adding 10 μM Forskolin (Fsk), every 24h for 72 h.

### RNA extraction

Total RNA extraction was performed using NucleoSpin RNA XS kit (Clontech Laboratories Inc. Cat. No. 740902). Briefly, cells were detached as mentioned in previous section, snap frozen in liquid nitrogen and stored at −80ᵒC overnight. Next day, RNA was prepared from a total of 168 samples across three conditions (C, M and F) according to the manufacturer’s instructions. RNA quality was assessed using High Sensitivity Fragment Analyzer.

### cDNA library preparation

cDNA library was prepared using SMART-Seq v4 3’ DE kit according to the manufacturer’s instructions (Clontech Laboratories Inc. Cat. No. 635040). For each clone, 1 ng of total RNA was used as input to synthesize cDNA which was subsequently amplified. Control RNA was provided with the kit. All samples were processed concurrently during library preparation. Prior to purification, the amplified cDNA’s were grouped in fourteen pools of twelve samples each with a unique barcode (Clonetech Oligo dT in-line indexes or IL 1-12) ligated to the 3’end during cDNA synthesis. Pooled and purified libraries were fragmented using Nextera XT DNA Sample Preparation Kit (Illumina, Cat. Nos. FC-131-1024, FC-131-1096). Illumina sequencing indexes (provided with the SMART-Seq v4 3’ DE kit) were added to the fragmented libraries and indexed libraries were amplified. For intermittent purification steps, Agencourt AMPure XP PCR purification kit (5 ml Beckman Coulter Part No. A63880; 60 ml Beckman Coulter Part No. A63881) was used. Purified samples were analyzed on a High Sensitivity Fragment Analyzer (Agilent Technologies). The pooling strategy and the combinations of Illumina indices employed in this experiment are given in Table 1.

### Denaturation and dilution of purified libraries

Purified libraries were denatured and diluted as outlined in the “Denature and Dilute Libraries Guide” designed for the NextSeq sequencing platform by Illumina. Libraries were sequenced across three different sequencing runs with final loading concentration equal to 1.8 pM. Un-indexed control library PhiX (NextSeq™ PhiX Control Kit FC-110-3002 by Illumina) was denatured and diluted to 1.8 pM according to the “Denature and Dilute Libraries Guide” and used as a spike-in to provide improved nucleotide diversity.

### Sequencing

The choice of sequencing kits and the decision on number of pools to be sequenced in a single run were based on the ENCODE guidelines for RNA-seq experiments. The libraries were sequenced on NextSeq 500 platform using NextSeq^®^ 500/550 High Output Kits v2 (75 cycles; Cat. No. FC-404-2005) for paired-end sequencing. In this, 35 read-1 cycles and 40 read-2 cycles were performed with adapter trimming feature turned off. Transcript/gene sequence information was exclusively present in read 1 whereas most of read 2 included sequence of the matched IL’s.

### Data acquisition and analysis

The FASTQ files obtained were de-multiplexed using SMART-Seq DE3 Demultiplexing Software by Clonetech, which organized reads into individual libraries by IL’s. Bowtie was used to align the reads to the human genome assembly hg19 [55]. The aligned reads were then processed by RSEM to generate un-normalized, TPM, and FPKM counts [56]. Sequencing run quality was analyzed and clones that did not have high-fidelity sequence data or that had low Sanger quality scores were eliminated from further analysis. Further review and inspection of the data set revealed no significant variation between the sequencing runs. Furthermore, upon distribution analysis, no skew was detected between runs. Clones that did not have sufficient count results in all three conditions, Control (C), MDI (M), and Forskolin (F), were disregarded from this portion of the study. In total, 12 out of the 168 sequenced samples were excluded from this portion of the study, leaving 52 clones under 3 conditions (C, M and F).

### Clustering

A matrix containing TPM counts of genes in “M” clones, and values for mean droplet sizes was analyzed using the Broad Institute “Morpheus” software (https://software.broadinstitute.org/morpheus/). Genes with a correlation coefficient greater than 0.5 and less than −0.5 (to account for negative monotonic trends) to lipid droplet size were then used for hierarchical clustering, using one minus the Spearman’s coefficient of correlation to calculate distances between clusters and then used the average of complete and single linkage to generate the dendrogram shown in Figure 2b.

### Pathway Analysis

Differential expression between clusters defined in Figure 2b was calculated using DESeq [57]. Lists of differentially expressed genes were uploaded to ToppGene [58] via the ToppFun uploader. ToppFun default parameters were used to generate KEGG and GO Term lists. Pathways were considered to be significantly enriched if the p-adj value was less than 0.05.

### Immunofluorescence

Antibodies against human adiponectin and leptin were purchased from Thermo Fisher (Adiponectin Monoclonal Antibody (19F1); Cat. No. MA1-054, Leptin Polyclonal Antibody; Cat. No. PA1-051). Cells were rinsed twice with 1X PBS (pH 7.4) and fixed with 4% PFA for 15 minutes followed by three washes with 1X PBS. Fixed cells were permeabilized using freshly prepared permeabilization buffer (1% FBS and 0.5% Triton X-100 in 1X PBS) for 30 minutes and incubated with a cocktail of anti-adiponectin (1:100) and anti-leptin (1:100) primary antibodies for 2 hrs.

Cells were washed three times with permeabilization buffer followed by incubation with a cocktail of anti-mouse (Alexa Fluor^®^ 594 Goat anti-mouse, Thermo Fisher A11005; 1:1000), anti-rabbit (Alexa Fluor^®^ 488 Donkey anti-rabbit, Thermo Fisher A21206; 1:1000) and DAPI (Hoechst 33342, trihydrochloride trihydrate; Thermo Fisher H3570; 1:2000) for 30 minutes. After three washes with permeabilization buffer, cells were mounted using ProLong™ Gold Antifade Mountant (Thermo Fisher P36930). Antibody dilutions were prepared fresh in permeabilization buffer. All incubation and wash steps were performed at room temperature on a shaker.

### Image acquisition and processing

Cells were imaged with a Zeiss Axiovert 200M Fluorescence microscope. ImageJ (FIJI) was used for image processing and normalization. Mean fluorescence intensity was calculated on individual cells and the values were used to plot frequency distribution using GraphPad Prism v.7. Adipocyte droplet size was quantified using Fiji software from brightfield images of individual wells of 384-well multiwells taken with a 20x objective. Image contrast was enhanced (background subtraction 50, enhance contrast 0.5), and images were converted to 8-bit greyscale images. The images were subsequently binarized, and size in pixels^2^was recorded for all objects in the field with a circularity between 0.5 and 1.

### Real Time-PCR

Total RNA was extracted and reverse-transcribed according to the manufacturer’s protocol (iScript™ cDNA Synthesis Kit, BIO-RAD). cDNA was loaded in duplicate and qPCRs were performed using the BIO-RAD CFX System (BIO-RAD, Hercules, CA, USA) and iQ™ SYBR Green™ Supermix (BIO-RAD cat. No. 1708882). Primers were designed through the Primer3 software platform. SYBR Primer specificity was confirmed via melt curve analysis.

### QUANTIFICATION AND STATISTICAL ANALYSIS

GraphPad Prism 7.0 was used for all analyses. Parametric or non-parametric test were chosen based on results from the D’Agostino-Pearson omnibus normality test, and are described in each figure. Heatmaps were plotted using Morpheus (Broad Institute).

---

**Supplementary Figure S2.**
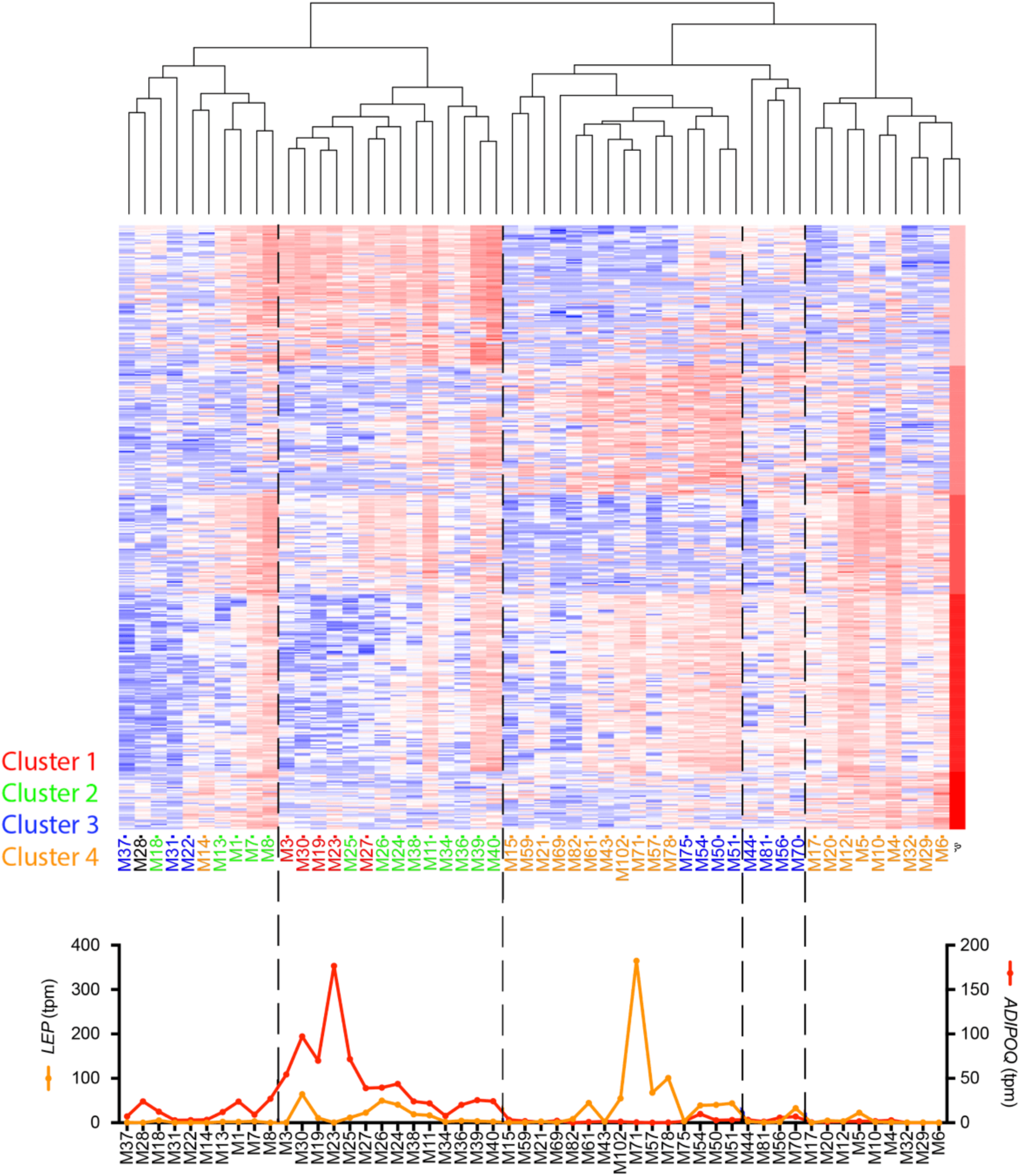
Alternative approach to hierarchical clustering. Unsupervised hierarchical clustering of TPM values for the 1500 genes displaying the highest variance in the M condition. K-means clustering was used to define gene clusters across the vertical axis. Clone identifiers are color coded to identify the cluster each clone associated with using the strategy depicted in Figure 2. Bottom panel shows TPM values for *LEP* and *ADIPOQ* across the defined clusters.

**Supplementary Table S1.**
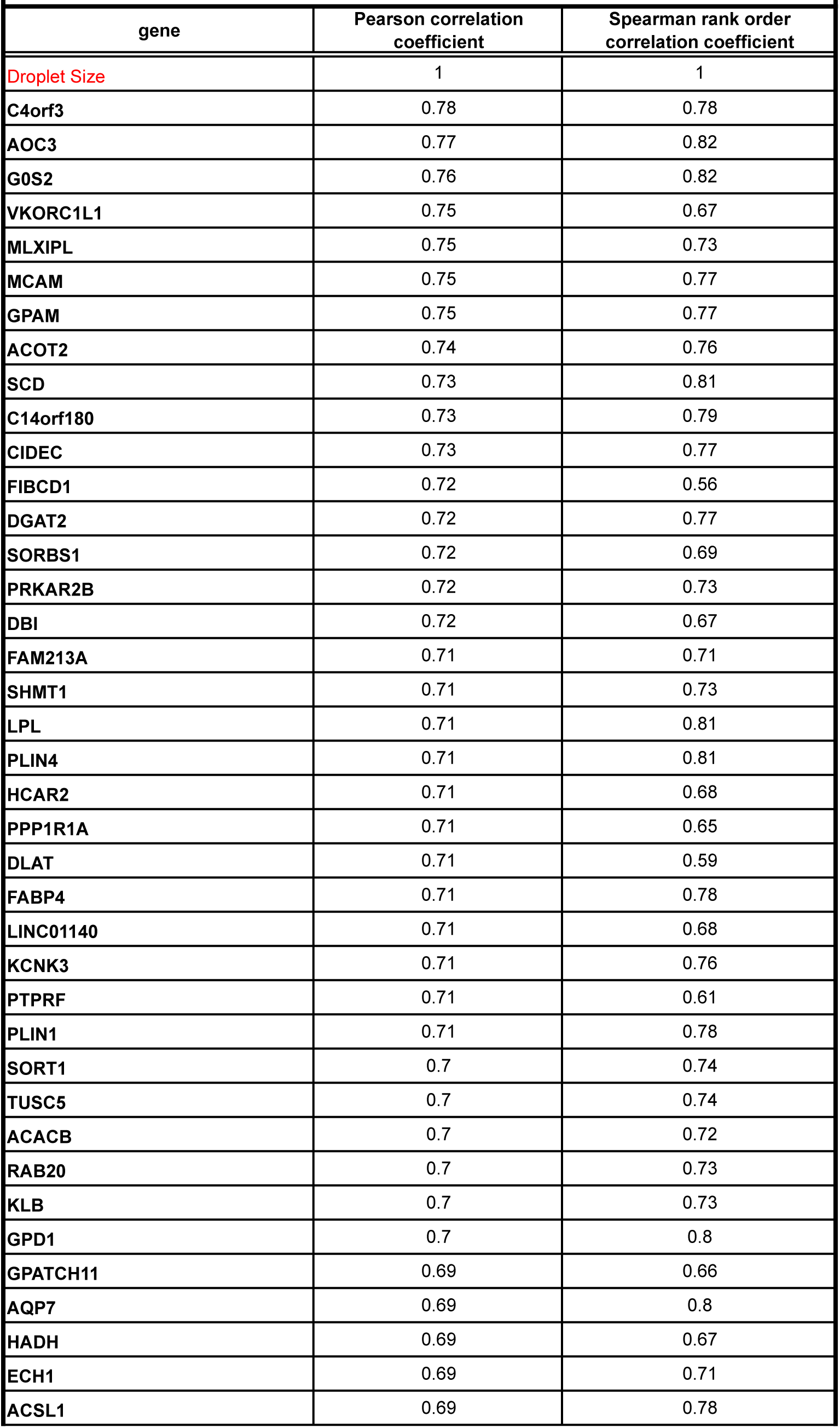
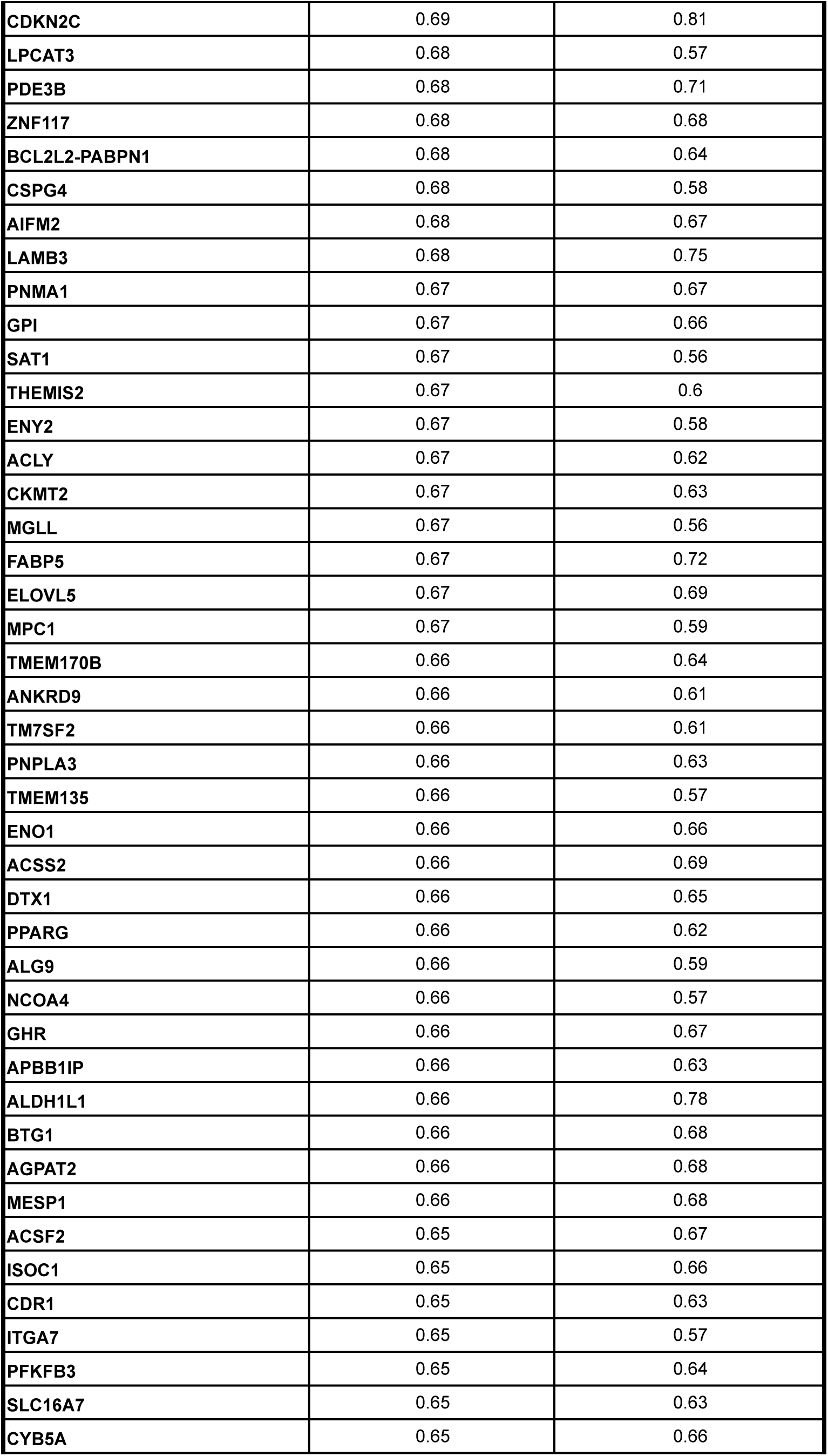
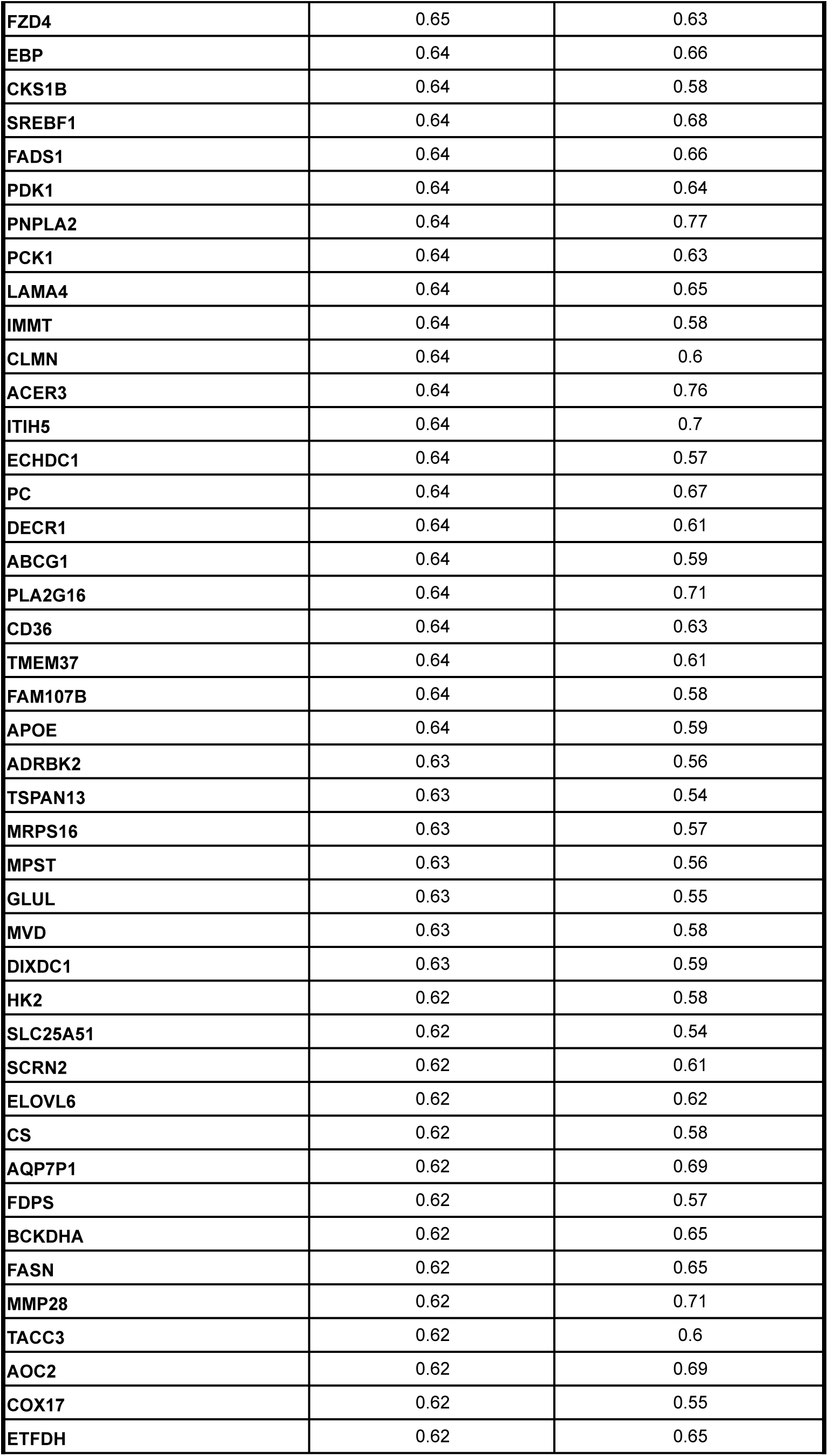
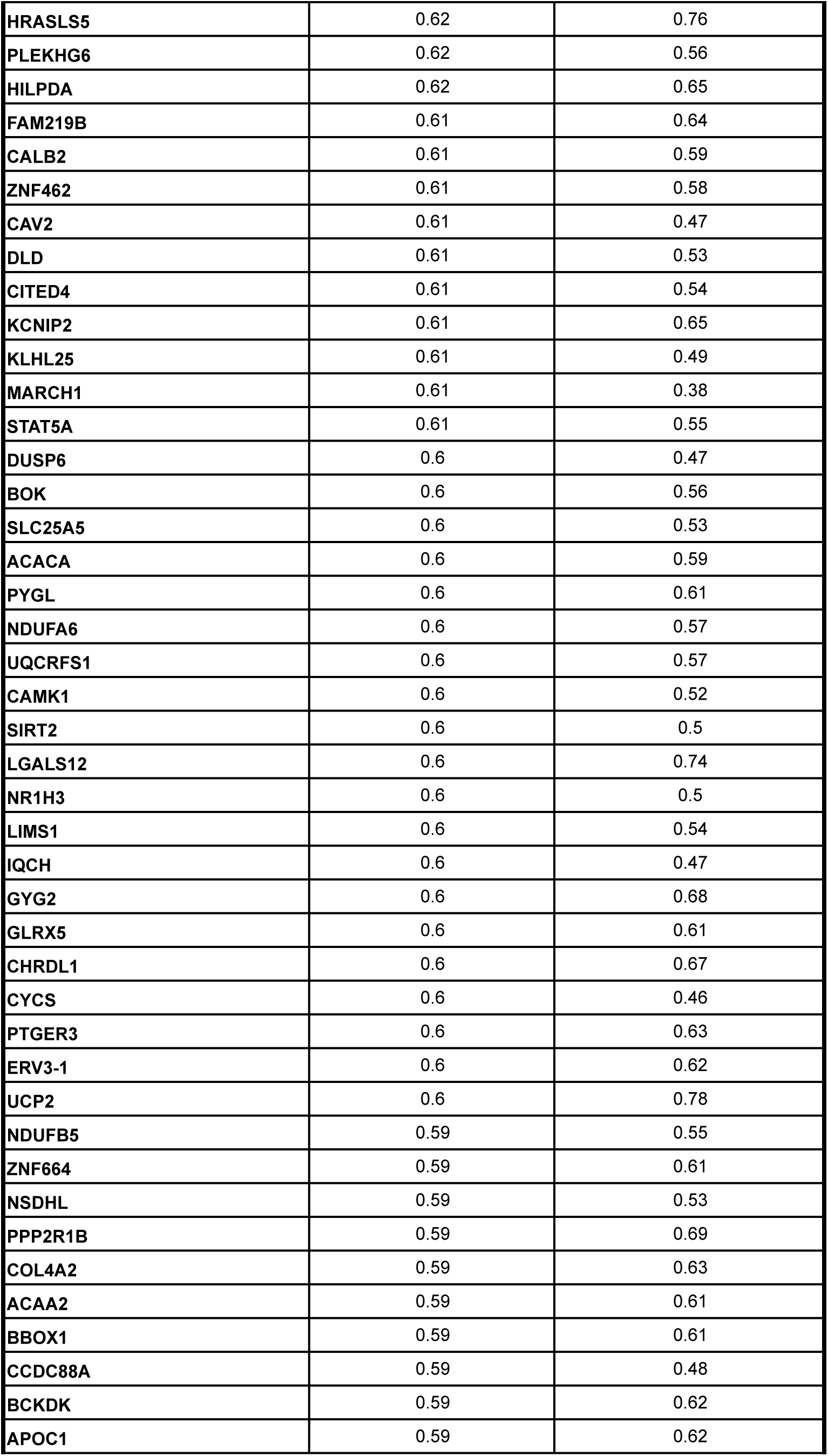
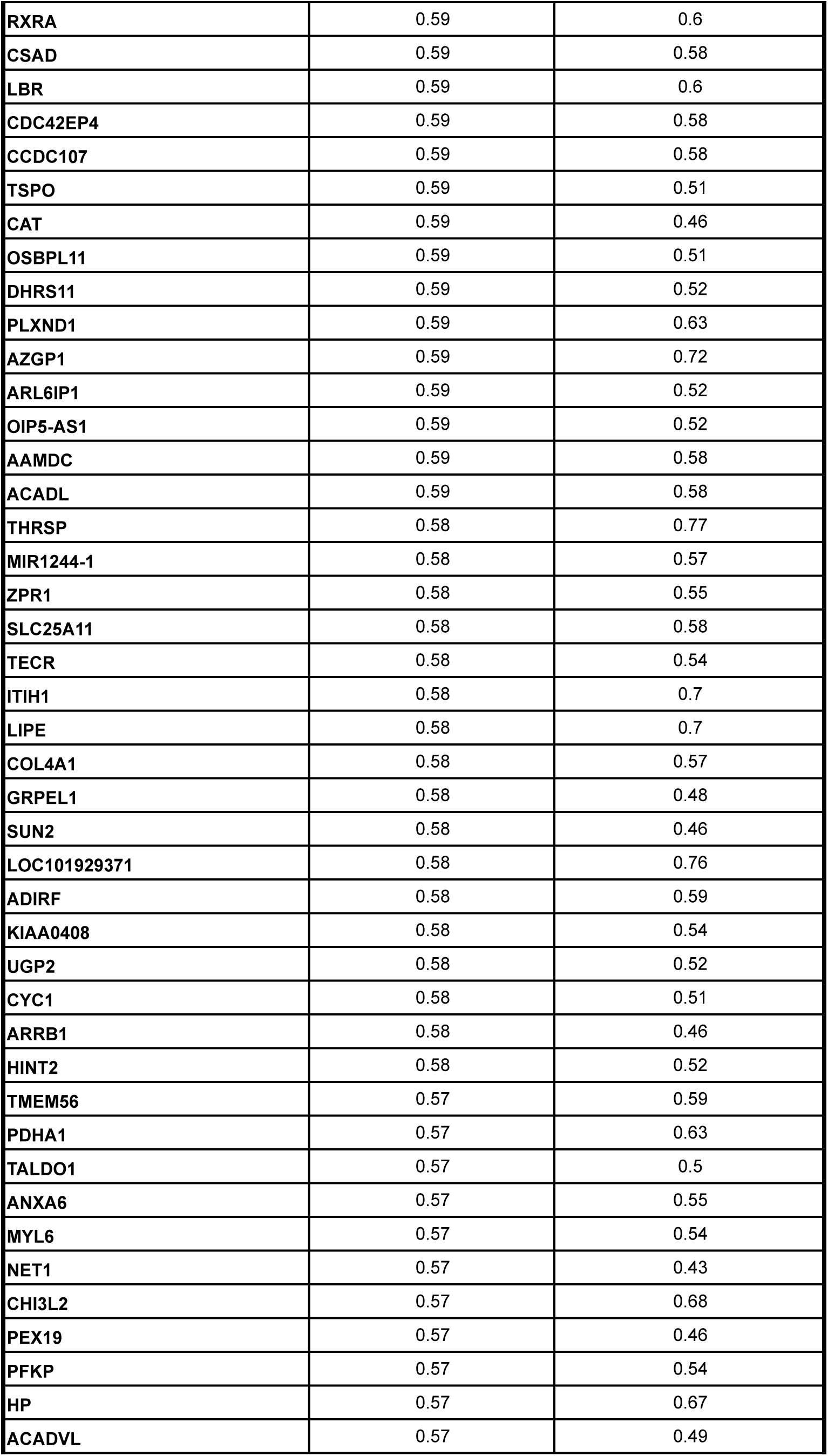
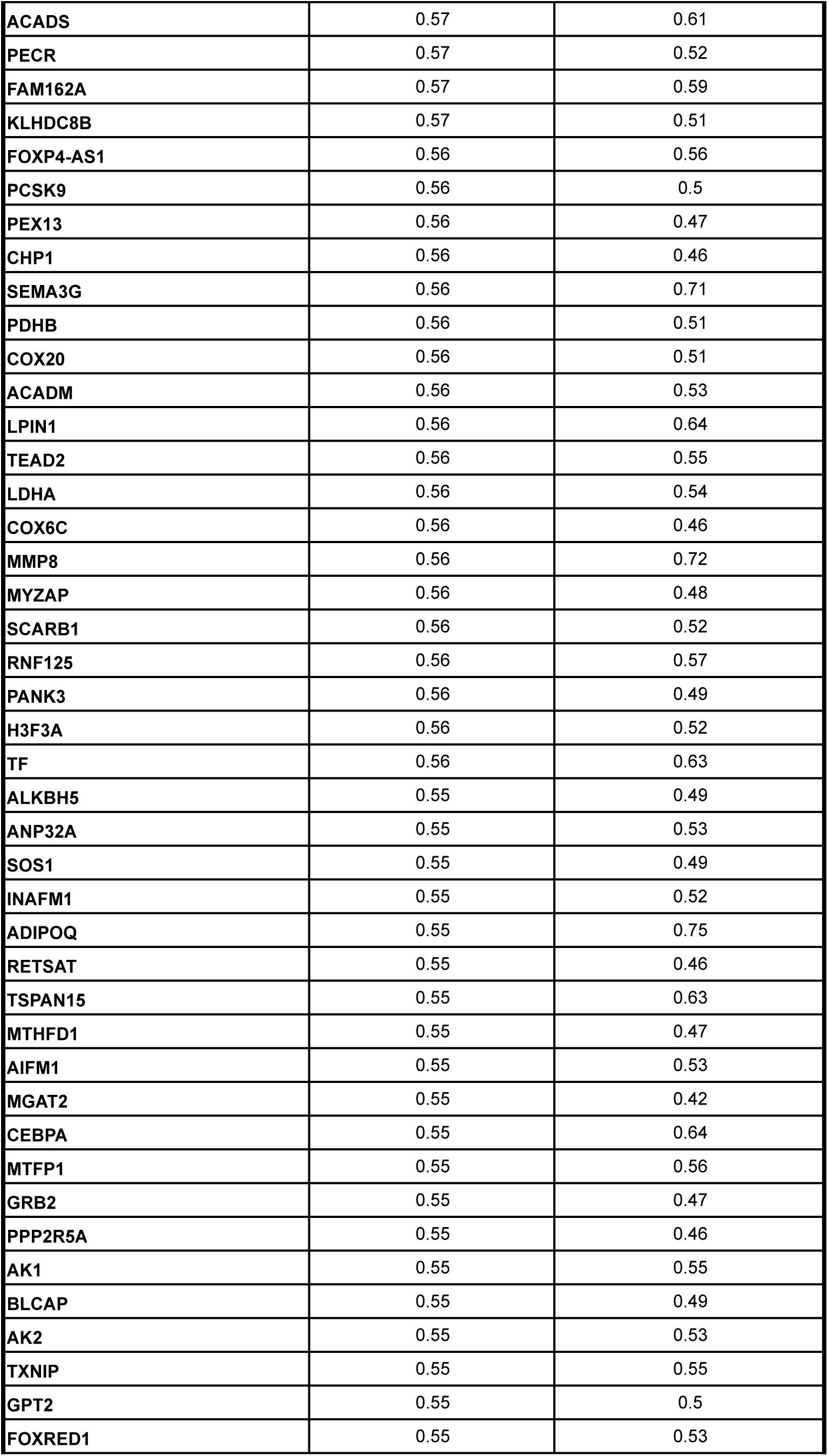
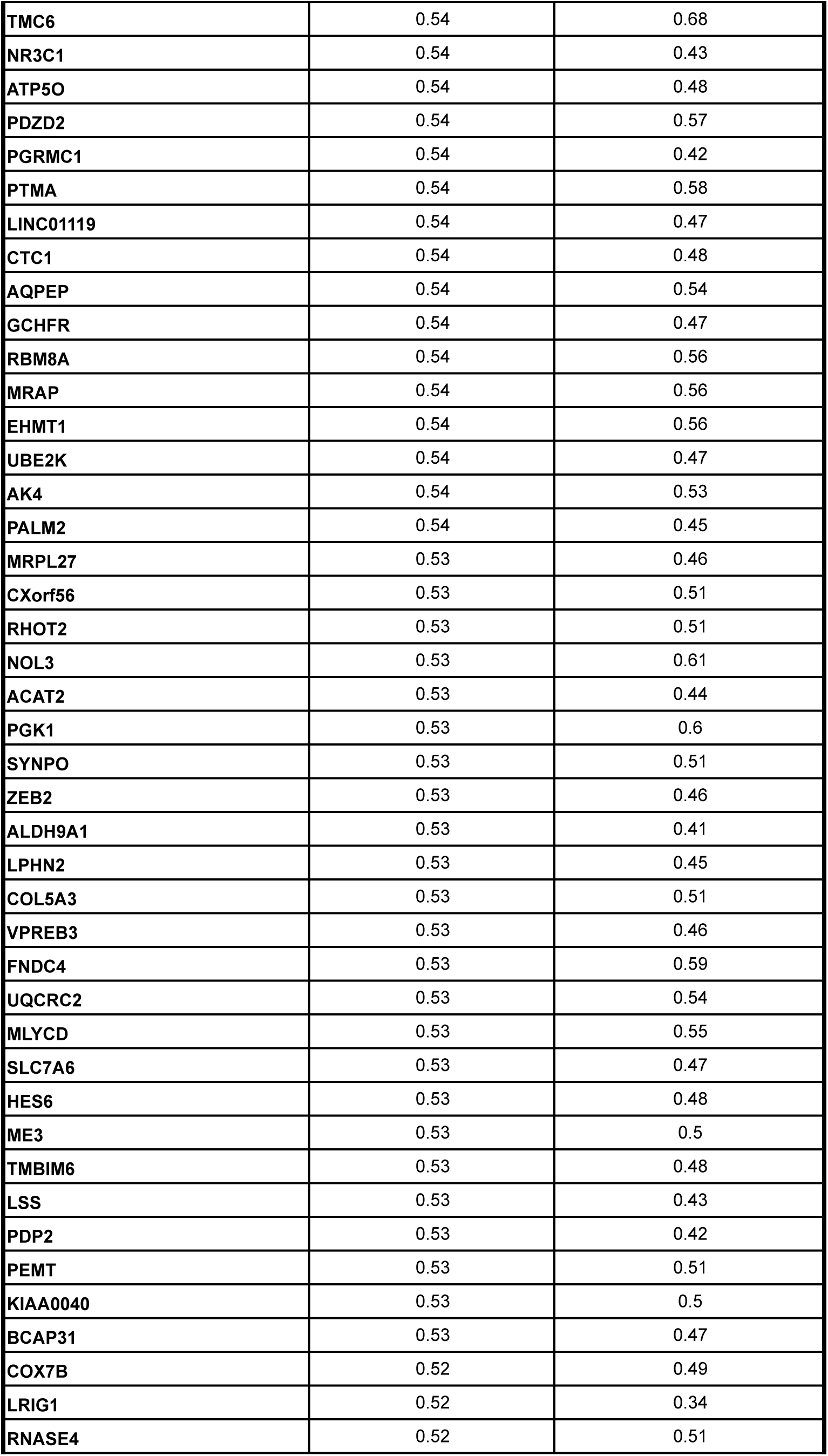
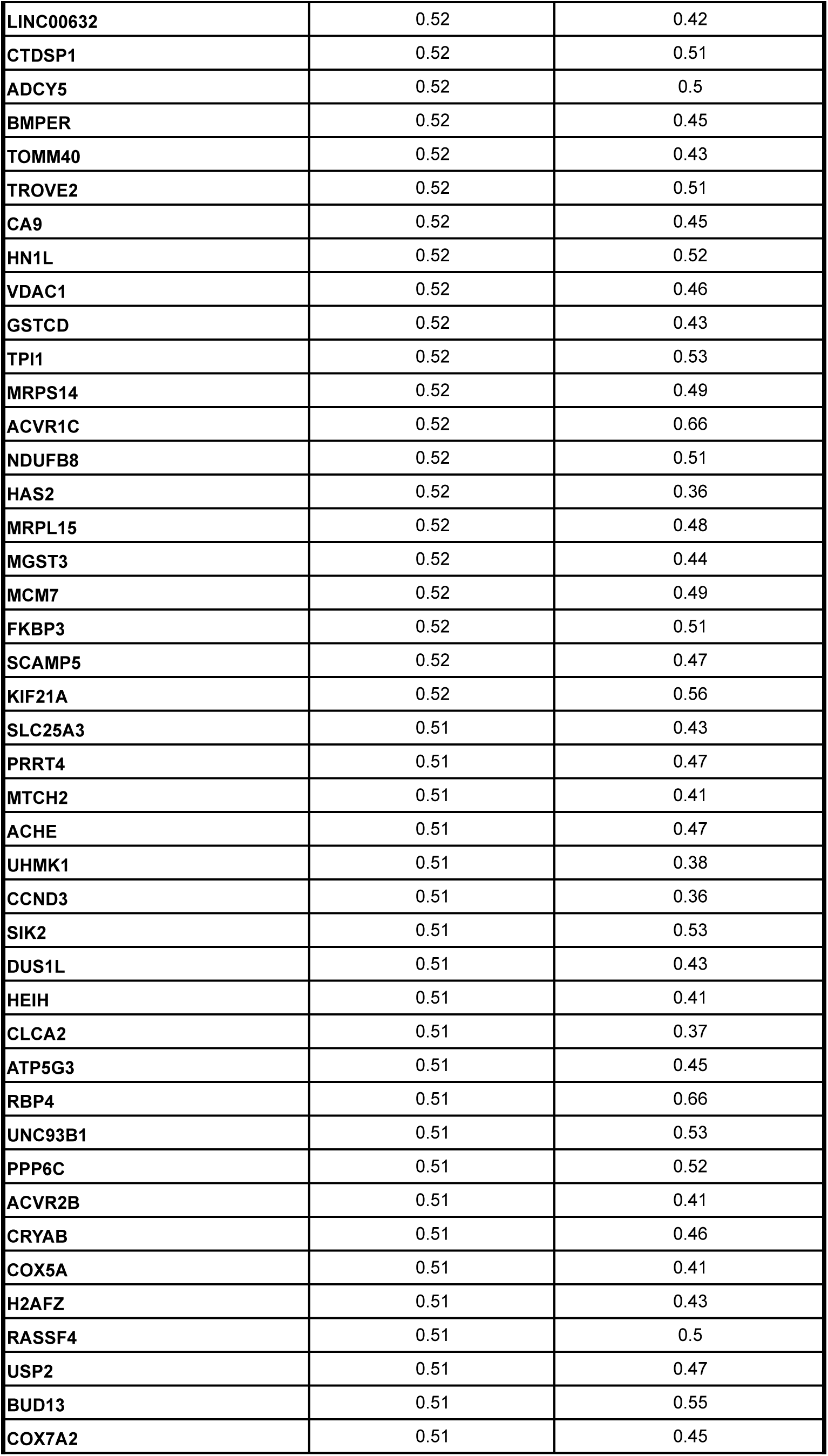
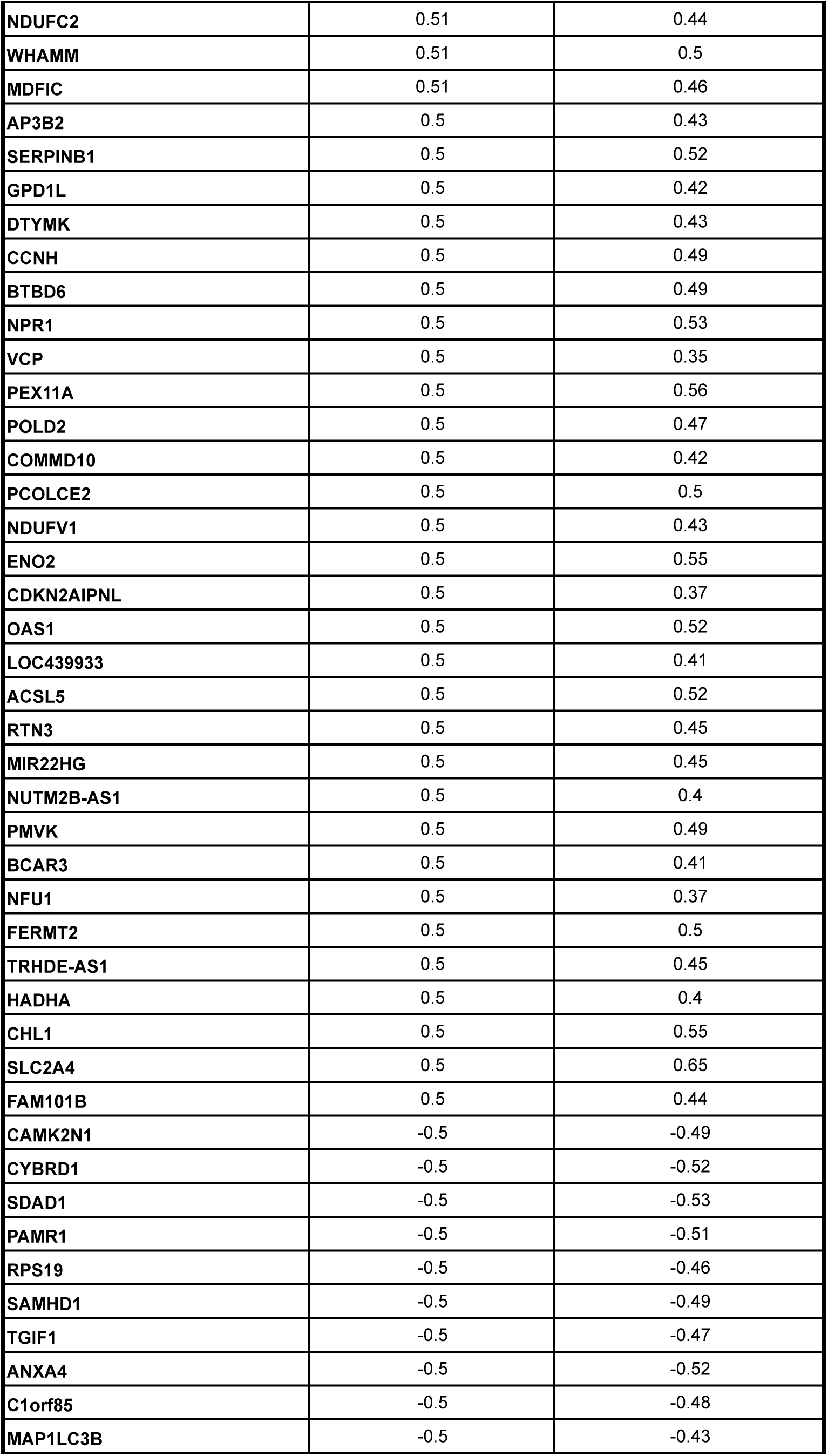
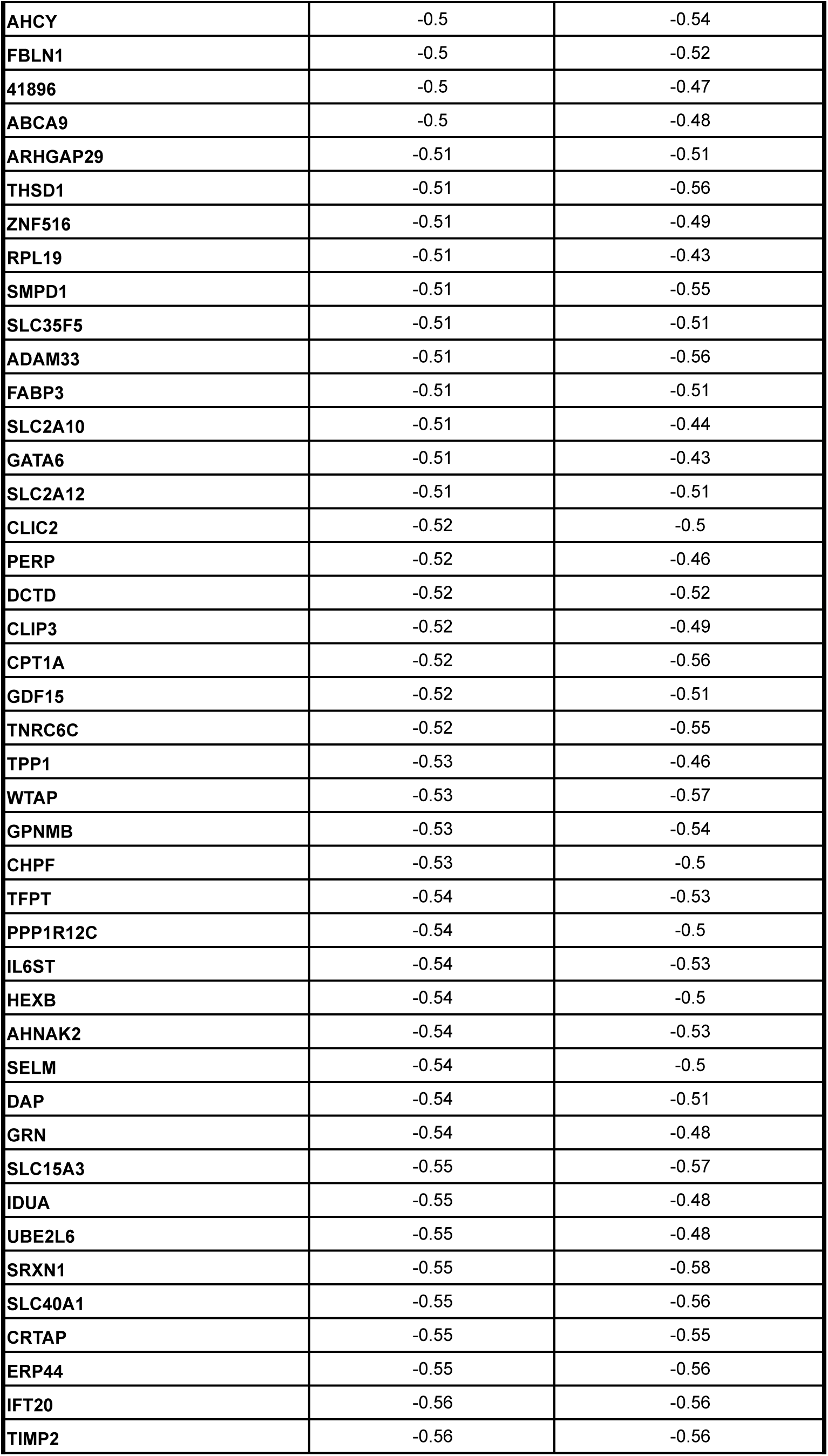
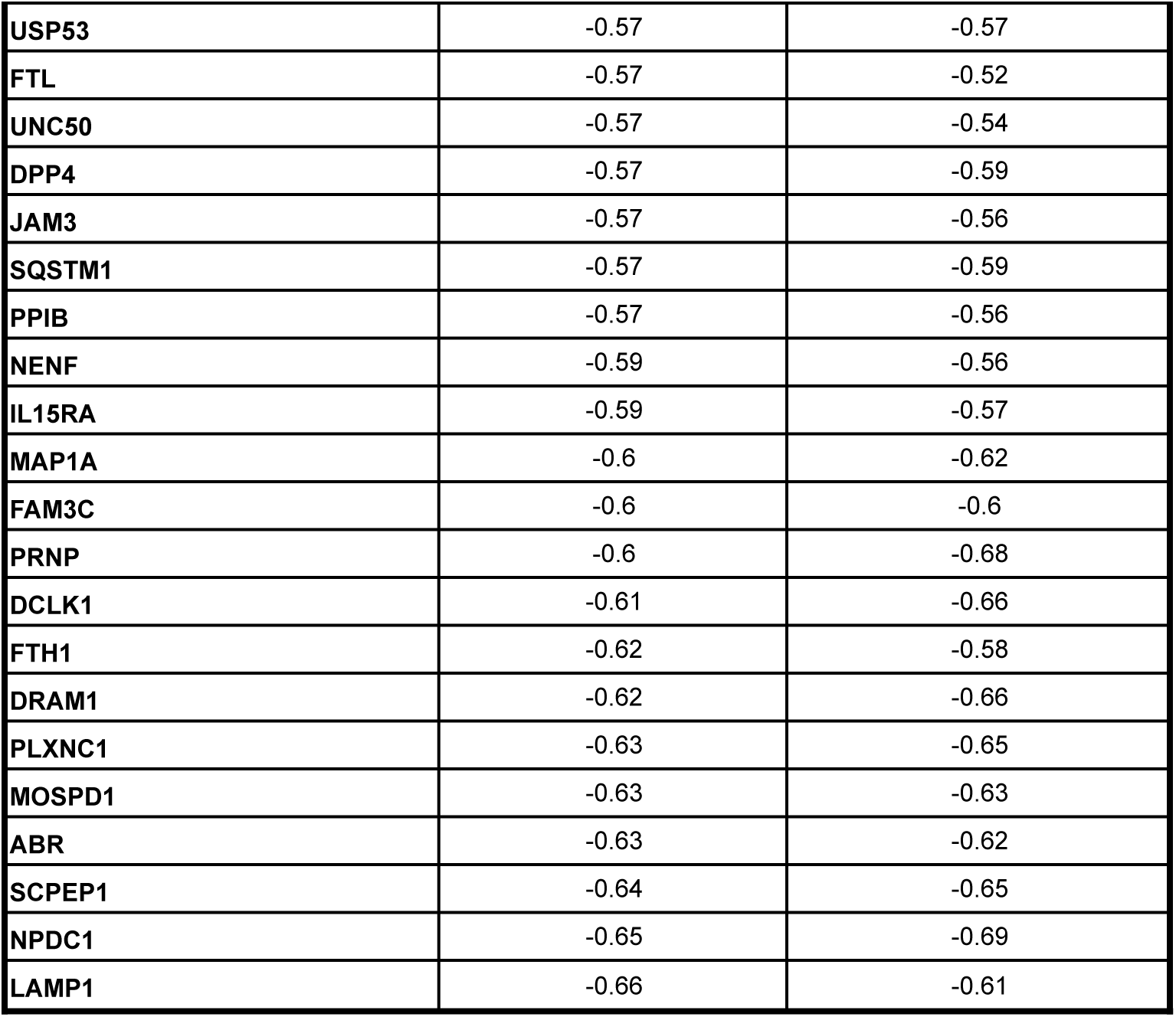
Genes correlated with lipid droplet size

**Supplementary Table S2.**
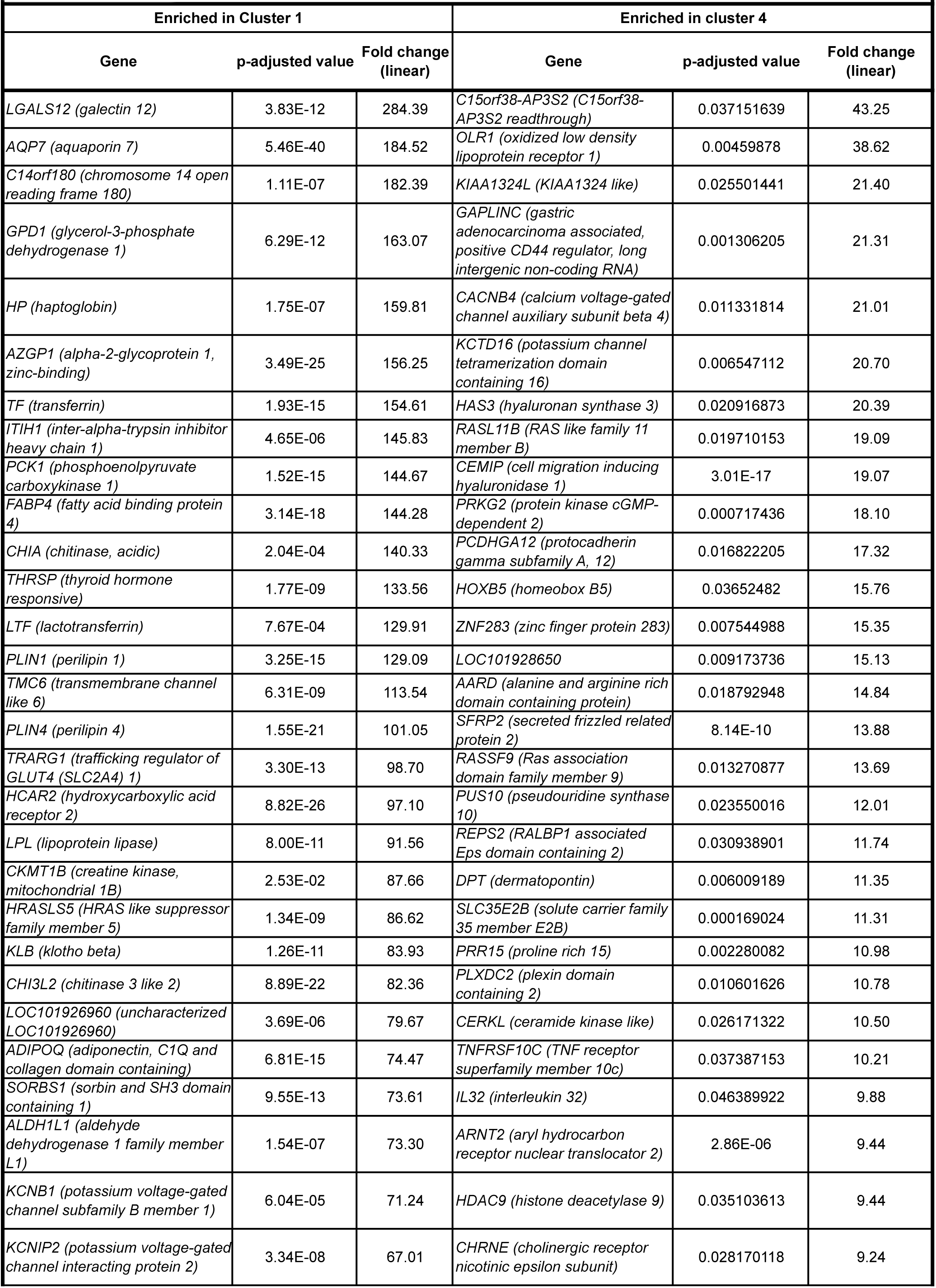
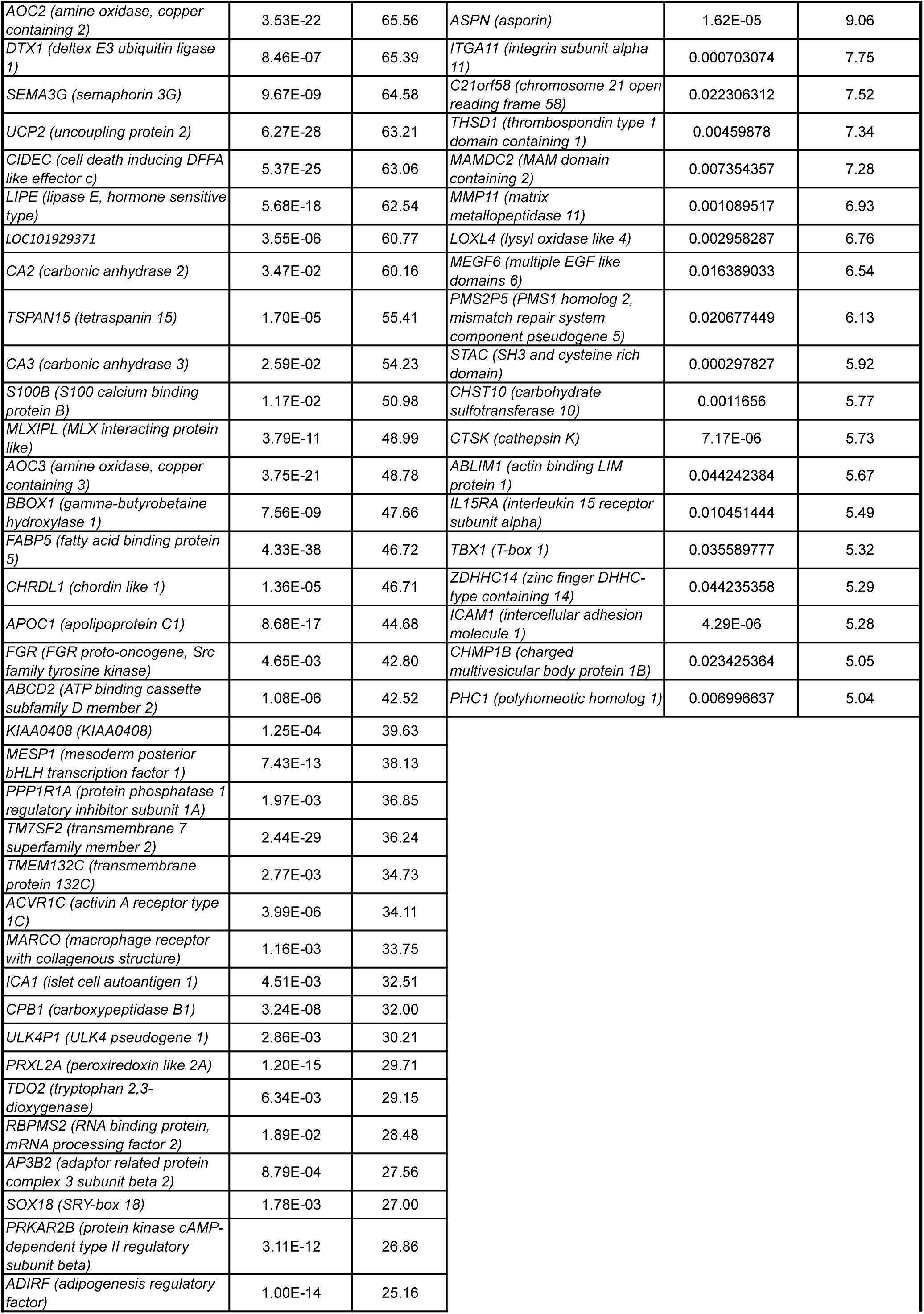
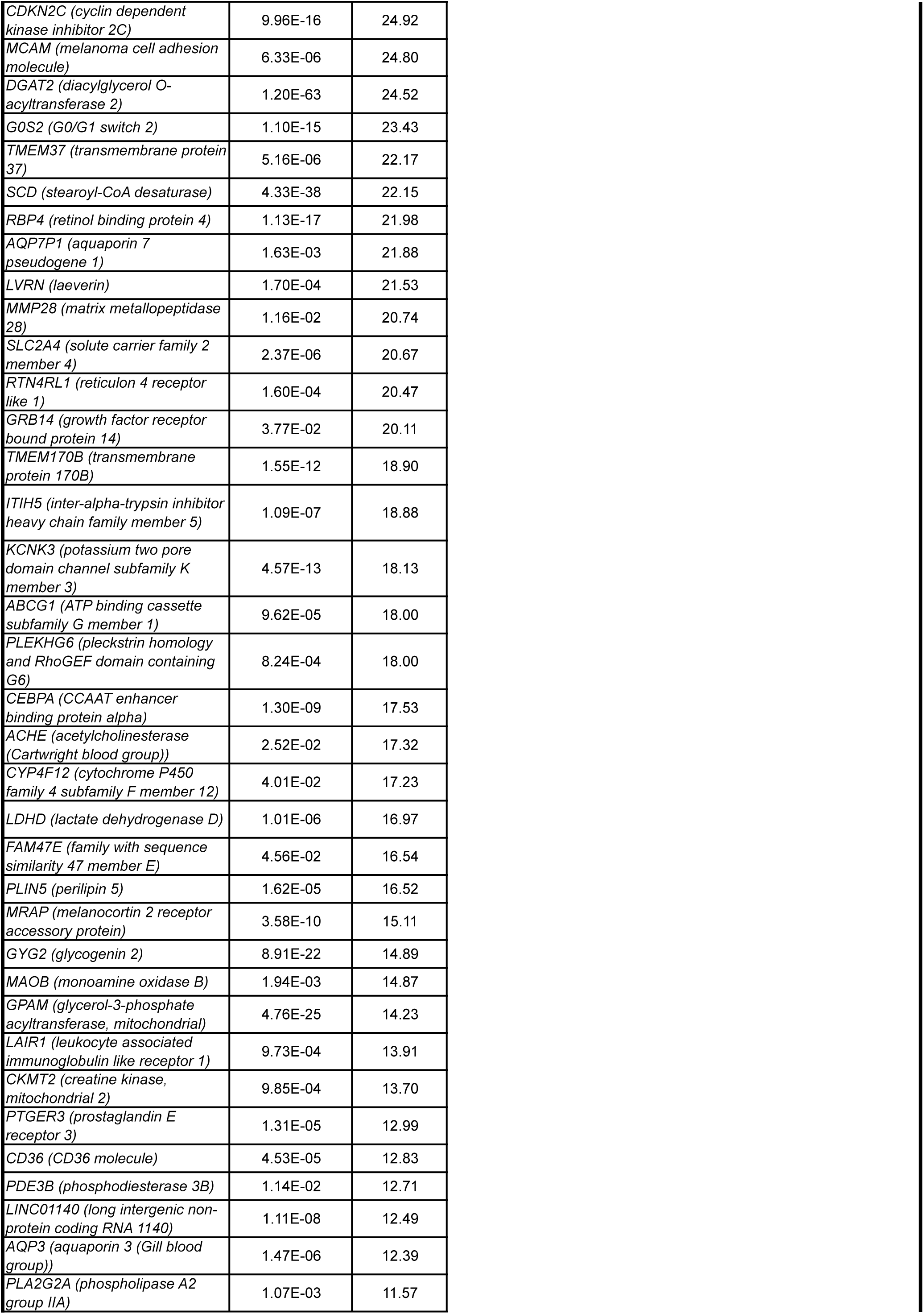
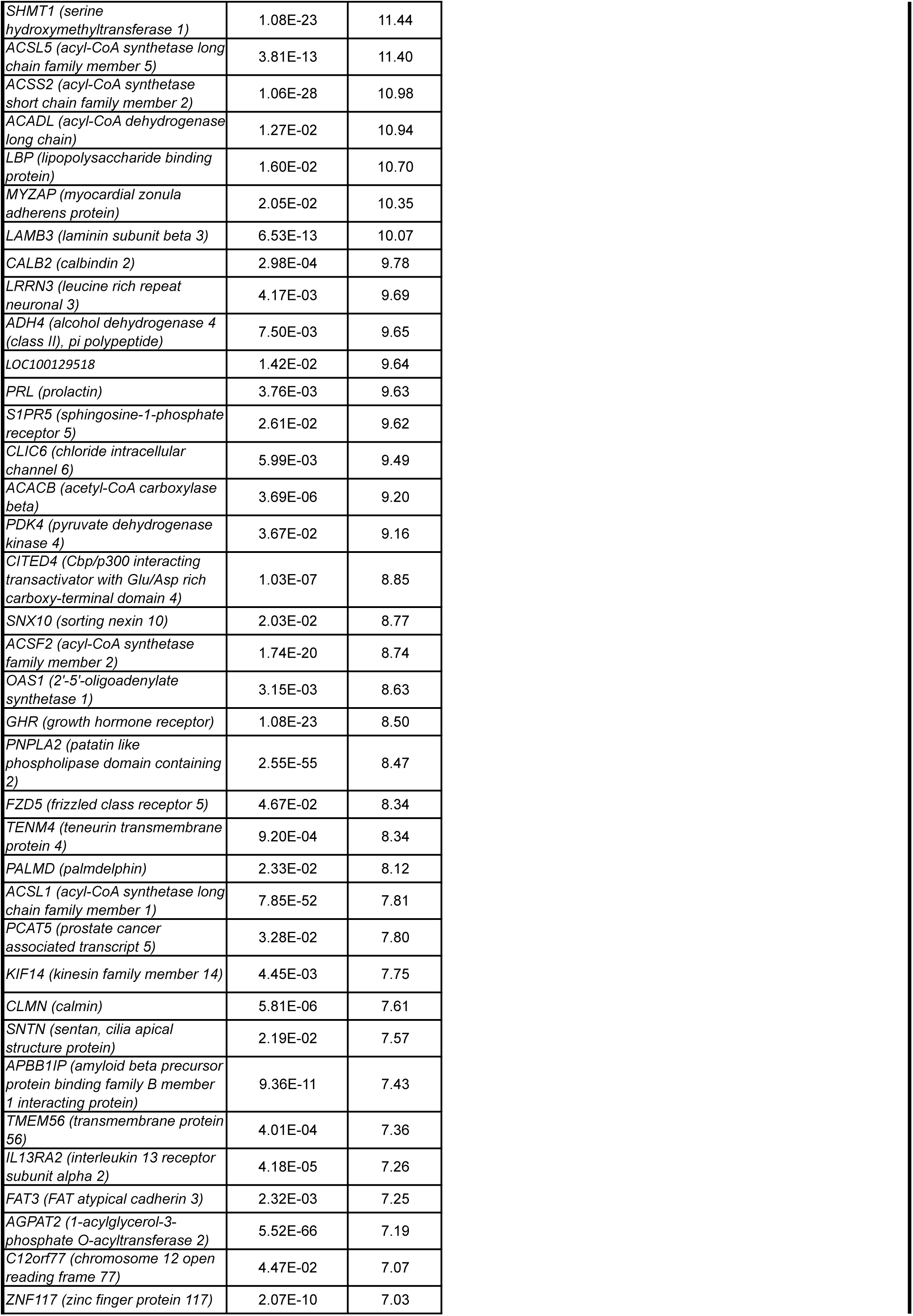
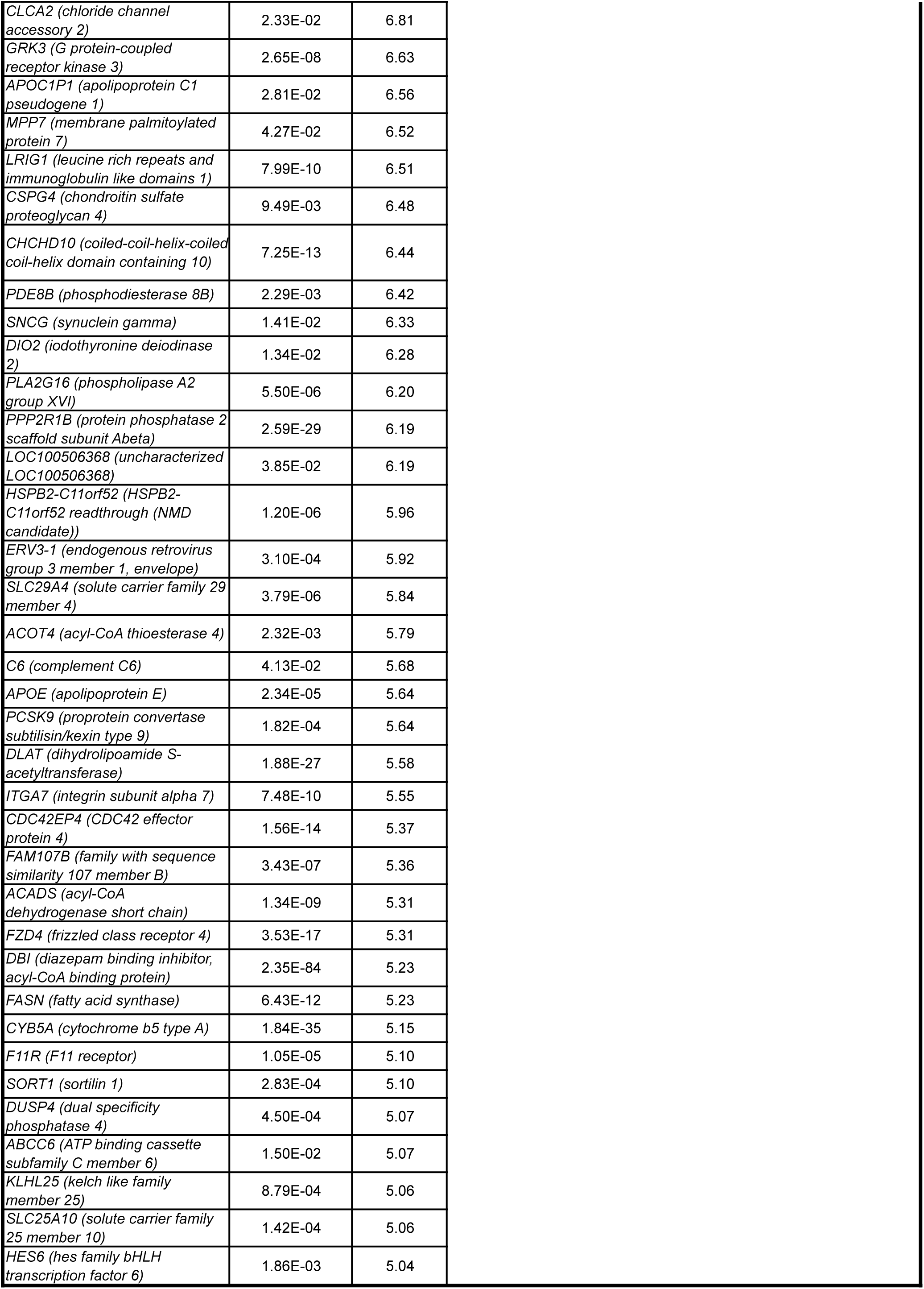
Genes differentially expressed between Cluster 1 and Cluster 4

**Supplementary Table S3.**
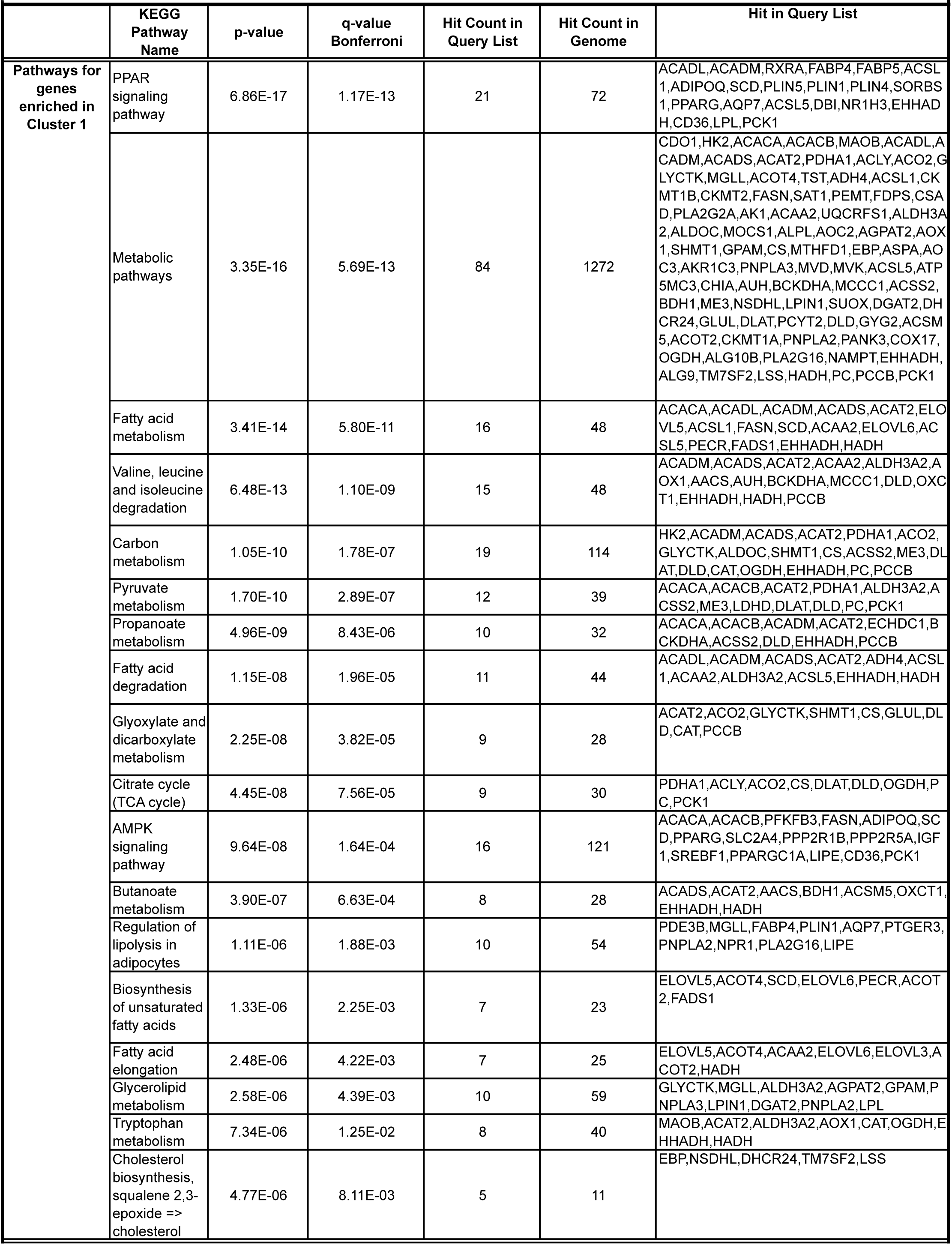
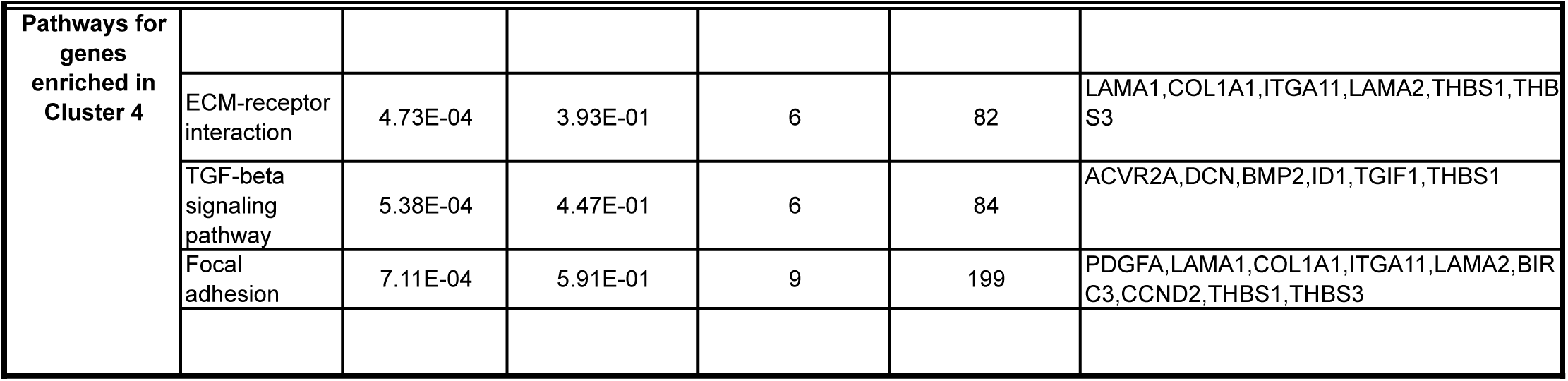
Pathway enrichement analysis of genes differentially expressed between Cluster 1 and Cluster 4.

**Supplementary Table S4.**
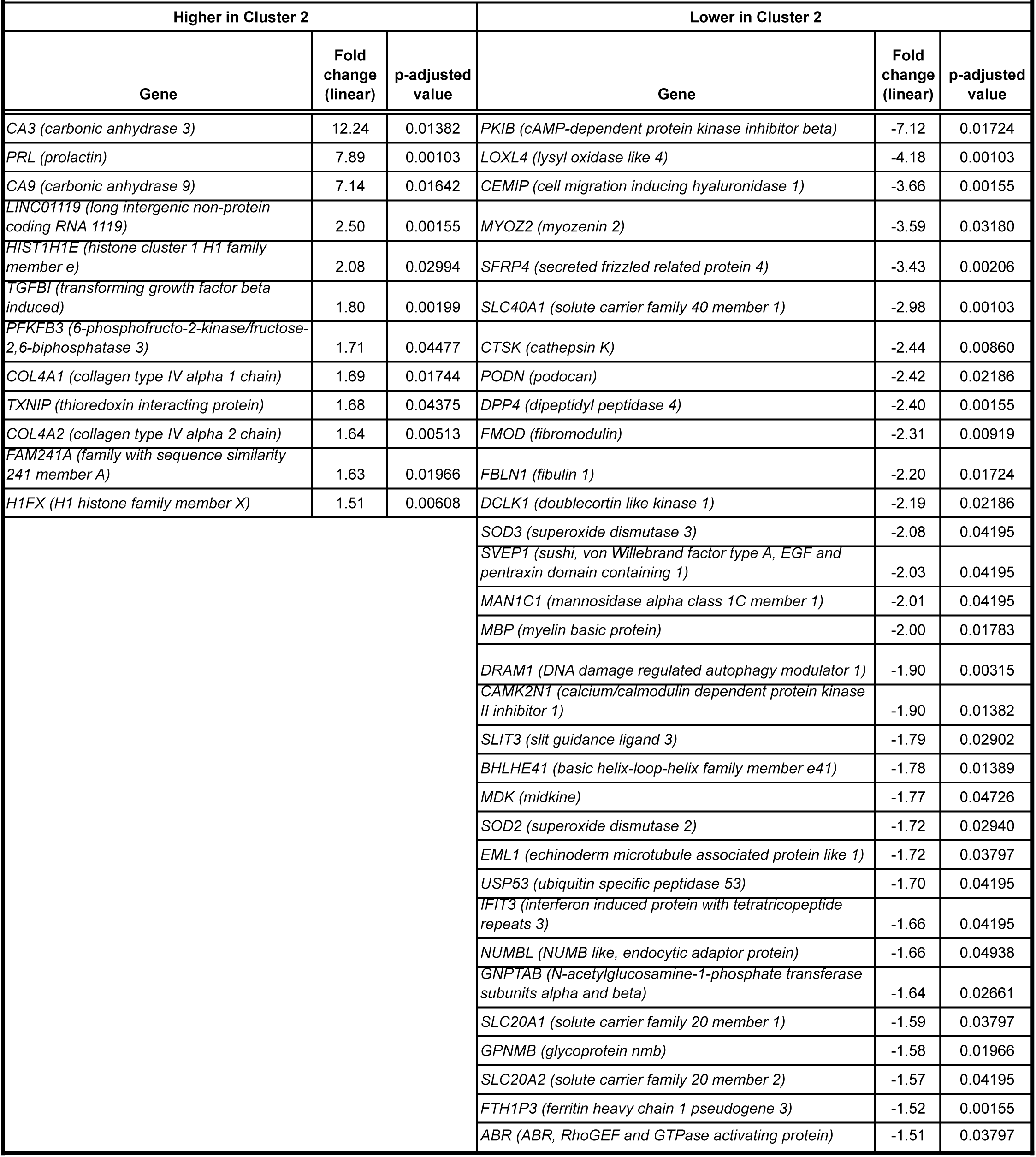
Genes differentially expressed between Cluster 2 and all others.

**Supplementary Table S5.** Pathway enrichment analysis of genes differentially expressed between Cluster 2 and all others.

|  | KEGG Pathway Name | p-value | q-value<br>Bonferroni | Hit Count<br>in Query<br>List | Hit<br>Count<br>in<br>Genom<br>e | Hit in Query List |
| --- | --- | --- | --- | --- | --- | --- |
| **Pathways for**<br>**genes enriched**<br>**in Cluster 2** | Glycolysis (Embden-Meyerhof pathway), glucose => pyruvate | 6.24E-16 | 2.55E-13 | 9 | 25 | GPI,GAPDH,PKM,PFKL,PFKP,ENO1,ENO2,PGK1,TPI1 |
|  | Glycolysis / Gluconeogenesis | 2.45E-13 | 1.00E-10 | 10 | 67 | GPI,GAPDH,PKM,PFKL,PFKP,LDHA,ENO1,ENO2,PGK1,TPI1 |
|  | Glycolysis, core module involving three-carbon compounds | 5.08E-12 | 2.08E-09 | 6 | 12 | GAPDH,PKM,ENO1,ENO2,PGK1,TPI1 |
|  | Biosynthesis of amino acids | 1.15E-09 | 4.69E-07 | 8 | 75 | GAPDH,PKM,PFKL,PFKP,ENO1,ENO2,PGK1,TPI1 |
|  | Carbon metabolism | 1.47E-09 | 6.01E-07 | 9 | 114 | GPI,GAPDH,PKM,PFKL,PFKP,ENO1,ENO2,PGK1,TPI1 |
|  | Gluconeogenesis, oxaloacetate => fructose-6P | 8.30E-09 | 3.40E-06 | 5 | 17 | GAPDH,ENO1,ENO2,PGK1,TPI1 |
|  | HIF-1 signaling pathway | 2.77E-07 | 1.13E-04 | 7 | 101 | GAPDH,PFKFB3,PFKL,LDHA,ENO1,ENO2,PGK1 |
|  | ECM-receptor interaction | 1.52E-06 | 6.22E-04 | 6 | 82 | COL4A1,COL4A2,COL6A1,COL6A2,LAMA4,ITGA5 |
|  | Central carbon metabolism in cancer | 9.44E-06 | 3.86E-03 | 5 | 65 | MYC,PKM,PFKL,PFKP,LDHA |
|  | Fructose and mannose metabolism | 1.27E-05 | 5.19E-03 | 4 | 33 | PFKFB3,PFKL,PFKP,TPI1 |
|  | Protein digestion and absorption | 4.62E-05 | 1.89E-02 | 5 | 90 | COL3A1,COL4A1,COL4A2,COL6A1,COL6A2 |
|  | PI3K-Akt signaling pathway | 1.18E-04 | 4.82E-02 | 8 | 342 | MYC,COL4A1,COL4A2,COL6A1,COL6A2,LAMA4,ITGA5,PRL |
|  | Focal adhesion | 2.34E-04 | 9.56E-02 | 6 | 199 | COL4A1,COL4A2,COL6A1,COL6A2,LAMA4,ITGA5 |
|  | Pentose phosphate pathway | 3.04E-04 | 1.25E-01 | 3 | 30 | GPI,PFKL,PFKP |
|  | RNA degradation | 3.63E-04 | 1.49E-01 | 4 | 77 | PFKL,PFKP,ENO1,ENO2 |
|  | Small cell lung cancer | 5.06E-04 | 2.07E-01 | 4 | 84 | MYC,COL4A1,COL4A2,LAMA4 |
|  | Amoebiasis | 8.38E-04 | 3.43E-01 | 4 | 96 | COL3A1,COL4A1,COL4A2,LAMA4 |
|  | Nitrogen metabolism | 2.50E-03 | 1.00E+00 | 2 | 17 | CA9,CA3 |
|  | Metabolic pathways | 3.02E-03 | 1.00E+00 | 13 | 1272 | GPI,P4HA2,MTMR6,GAPDH,PKM,NMRK1,PFKL,PFKP,LDHA,ENO1,ENO2,PGK1,TPI1 |
|  | Galactose metabolism | 8.21E-03 | 1.00E+00 | 2 | 31 | PFKL,PFKP |
|  | AGE-RAGE signaling pathway in diabetic complications | 9.49E-03 | 1.00E+00 | 3 | 99 | COL3A1,COL4A1,COL4A2 |
| **Pathways for**<br>**genes decreased**<br>**in Cluster 2** | Ferroptosis | 5.86E-08 | 2.77E-05 | 6 | 40 | GCLM,FTH1,FTL,MAP1LC3B,SLC40A1,PRNP |
|  | Mineral absorption | 1.49E-04 | 7.05E-02 | 4 | 51 | CYBRD1,FTH1,FTL,SLC40A1 |

**Supplementary Table S6.** Genes differentially expressed between progenitors for Cluster 1 and all others.

| Enriched in progenitors giving rise to Cluster 1 |  |  | Decreased in progenitors giving rise to Cluster 1 |  |  |
| --- | --- | --- | --- | --- | --- |
| Gene | Fold Change (Linear) | p adjusted value | Gene | Fold Change (Linear) | p adjusted value |
| *EVI2A* (ecotropic viral integration site 2A) | 3.81 | 0.033427884 | *NEFM* (neurofilament medium) | -174.53 | 0.022725442 |
| *CCDC69* (coiled-coil domain containing 69) | 2.53 | 0.035515468 | *PLSCR3* (phospholipid scramblase 3) | -43.61 | 0.00014831 |
|  |  |  | *SLC22A3* (solute carrier family 22 member 3) | -21.10 | 0.016915217 |
|  |  |  | *CXCL1* (C-X-C motif chemokine ligand 1) | -20.64 | 0.030480859 |
|  |  |  | *CXCL6* (C-X-C motif chemokine ligand 6) | -20.30 | 0.020206449 |
|  |  |  | *NBL1* (NBL1, DAN family BMP antagonist) | -16.66 | 0.042373531 |
|  |  |  | *MMP1* (matrix metalloproteinase 1) | -15.30 | 0.017375974 |
|  |  |  | *CXCL5* (C-X-C motif chemokine ligand 5) | -14.62 | 0.032115504 |
|  |  |  | *STC1* (stanniocalcin 1) | -8.13 | 0.030480859 |
|  |  |  | *INHBA* (inhibin subunit beta A) | -8.09 | 0.016915217 |
|  |  |  | *ID1* (inhibitor of DNA binding 1, HLH protein) | -7.12 | 0.047758672 |
|  |  |  | *DACT1* (dishevelled binding antagonist of beta catenin 1) | -6.94 | 0.000397007 |
|  |  |  | *ATOH8* (atonal bHLH transcription factor 8) | -6.30 | 0.035515468 |
|  |  |  | *JAM2* (junctional adhesion molecule 2) | -6.08 | 0.038224756 |
|  |  |  | *SCHIP1* (schwannomin interacting protein 1) | -5.95 | 0.00014831 |
|  |  |  | *ENC1* (ectodermal-neural cortex 1) | -4.92 | 0.018319432 |
|  |  |  | *ANXA2P3* (annexin A2 pseudogene 3) | -4.32 | 0.016377587 |
|  |  |  | *UBASH3B* (ubiquitin associated and SH3 domain containing B) | -4.05 | 0.024168171 |
|  |  |  | *PPIAL4G* (peptidylprolyl isomerase A like 4G) | -3.30 | 0.018319432 |
|  |  |  | *C1QTNF1* (C1q and TNF related 1) | -3.11 | 0.029872169 |
|  |  |  | *SLC2A1* (solute carrier family 2 member 1) | -3.11 | 0.016377587 |
|  |  |  | *SLC16A3* (solute carrier family 16 member 3) | -2.87 | 0.016377587 |
|  |  |  | *MIR210HG* (MIR210 host gene) | -2.79 | 0.007867149 |
|  |  |  | *RORA* (RAR related orphan receptor A) | -2.77 | 0.03479213 |
|  |  |  | *TPM1* (tropomyosin 1) | -2.70 | 0.039134788 |
|  |  |  | *SNHG9* (small nucleolar RNA host gene 9) | -2.46 | 0.03479213 |
|  |  |  | *TNS1* (tensin 1) | -2.38 | 0.031506188 |
|  |  |  | *CCDC124* (coiled-coil domain containing 124) | -2.36 | 0.042373531 |
|  |  |  | *FTH1P3* (ferritin heavy chain 1 pseudogene 3) | -2.30 | 0.033486735 |
|  |  |  | *CDKN2A* (cyclin dependent kinase inhibitor 2A) | -2.24 | 0.045705177 |
|  |  |  | *CLEC11A* (C-type lectin domain containing 11A) | -2.19 | 0.033317835 |
|  |  |  | *MICAL2* (microtubule associated monooxygenase, calponin and LIM domain containing 2) | -2.18 | 0.018319432 |
|  |  |  | *TNFRSF12A* (TNF receptor superfamily member 12A) | -2.15 | 0.018319432 |
|  |  |  | *FGF2* (fibroblast growth factor 2) | -2.00 | 0.049391853 |

**Supplementary Table S7.**
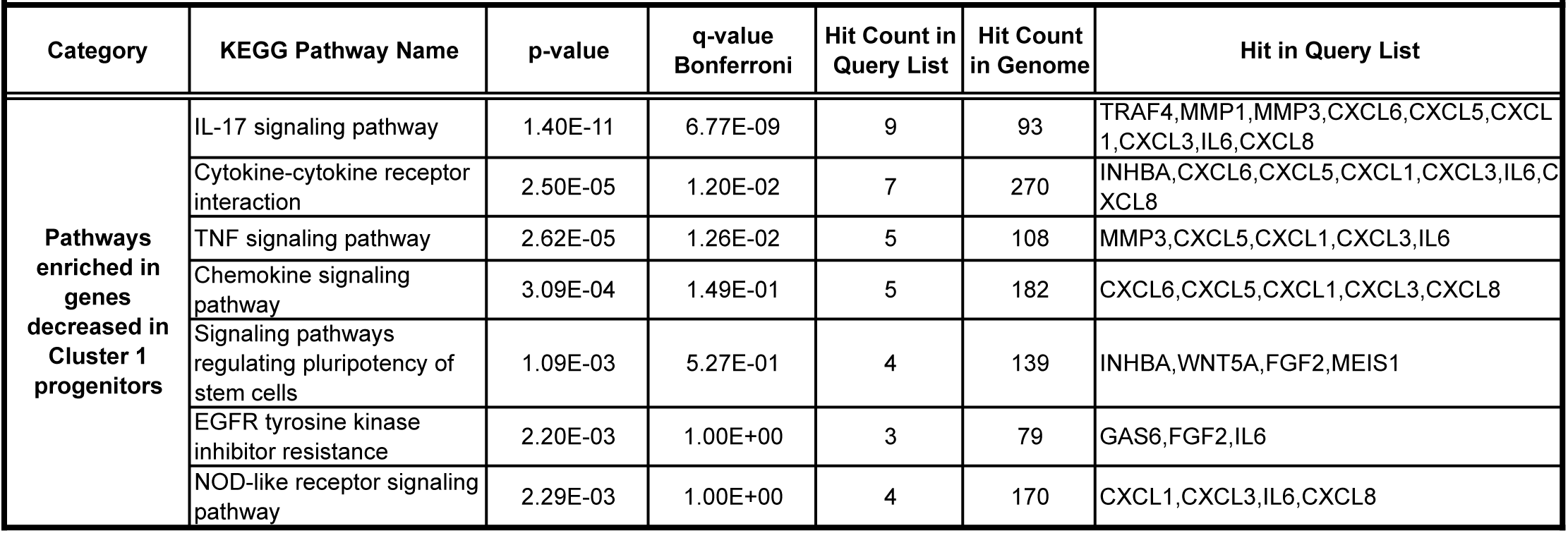
Pathway enrichment analysis of genes differentially expressed in progenitors for Cluster 1.

| Category | KEGG Pathway Name | p-value | q-value Bonferroni | Hit Count in Query List | Hit Count in Genome | Hit in Query List |
| --- | --- | --- | --- | --- | --- | --- |
| **Pathways enriched in genes decreased in Cluster 1 progenitors** | IL-17 signaling pathway | 1.40E-11 | 6.77E-09 | 9 | 93 | TRAF4,MMP1,MMP3,CXCL6,CXCL5,CXCL1,CXCL3,IL6,CXCL8 |
|  | Cytokine-cytokine receptor interaction | 2.50E-05 | 1.20E-02 | 7 | 270 | INHBA,CXCL6,CXCL5,CXCL1,CXCL3,IL6,CXCL8 |
|  | TNF signaling pathway | 2.62E-05 | 1.26E-02 | 5 | 108 | MMP3,CXCL5,CXCL1,CXCL3,IL6 |
|  | Chemokine signaling pathway | 3.09E-04 | 1.49E-01 | 5 | 182 | CXCL6,CXCL5,CXCL1,CXCL3,CXCL8 |
|  | Signaling pathways regulating pluripotency of stem cells | 1.09E-03 | 5.27E-01 | 4 | 139 | INHBA,WNT5A,FGF2,MEIS1 |
|  | EGFR tyrosine kinase inhibitor resistance | 2.20E-03 | 1.00E+00 | 3 | 79 | GAS6,FGF2,IL6 |
|  | NOD-like receptor signaling pathway | 2.29E-03 | 1.00E+00 | 4 | 170 | CXCL1,CXCL3,IL6,CXCL8 |

**Supplementary Table S8.** Genes differentially expressed between progenitors for Cluster 2 and all others.

| Enriched in progenitors giving rise to Cluster 2 |  |  | Decreased in progenitors giving rise to Cluster 2 |  |  |
| --- | --- | --- | --- | --- | --- |
| Gene | Fold Change<br>(Linear) | p adjusted | Gene | Fold Change<br>(Linear) | p adjusted |
| *CSF3* (colony stimulating factor 3) | 23.48 | 0.01673 | *ACTC1* (actin, alpha, cardiac muscle 1) | -11.39 | 0.03611 |
| *IL11* (interleukin 11) | 12.32 | 0.02760 | *PPP1R14C* (protein phosphatase 1 regulatory inhibitor subunit 14C) | -10.80 | 0.02929 |
| *CXCL8* (C-X-C motif chemokine ligand 8) | 11.25 | 0.00010 | *SPIN2A* (spindlin family member 2A) | -5.89 | 0.02614 |
| *OAS1* (2'-5'-oligoadenylate synthetase 1) | 10.57 | 0.00448 | *DLX2* (distal-less homeobox 2) | -5.54 | 0.02760 |
| *NR4A2* (nuclear receptor subfamily 4 group A member 2) | 8.15 | 0.00011 | *CLCA2* (chloride channel accessory 2) | -5.26 | 0.00878 |
| *IL1B* (interleukin 1 beta) | 8.08 | 0.02028 | *VAMP8* (vesicle associated membrane protein 8) | -5.01 | 0.02760 |
| *IFI6* (interferon alpha inducible protein 6) | 5.27 | 0.00000 | *ID1* (inhibitor of DNA binding 1, HLH protein) | -4.39 | 0.00878 |
| *LIF* (LIF, interleukin 6 family cytokine) | 4.09 | 0.02760 | *RELN* (reelin) | -3.91 | 0.01012 |
| *PTGS2* (prostaglandin-endoperoxide synthase 2) | 3.99 | 0.00290 | *CLEC3B* (C-type lectin domain family 3 member B) | -3.68 | 0.03078 |
| *CASC15* (cancer susceptibility 15) | 3.29 | 0.01337 | *THBD* (thrombomodulin) | -3.59 | 0.01012 |
| *IFIT1* (interferon induced protein with tetratricopeptide repeats 1) | 2.62 | 0.00912 | *MRAP2* (melanocortin 2 receptor accessory protein 2) | -3.13 | 0.04151 |
| *IRF7* (interferon regulatory factor 7) | 2.42 | 0.04085 | *HSPB2-C11orf52* (HSPB2-C11orf52 readthrough (NMD candidate)) | -3.04 | 0.01817 |
| *TMEM132A* (transmembrane protein 132A) | 2.42 | 0.03984 | *NFASC* (neurofascin) | -2.93 | 0.01337 |
| *PID1* (phosphotyrosine interaction domain containing 1) | 2.30 | 0.04306 | *HSPB7* (heat shock protein family B (small) member 7) | -2.57 | 0.04135 |
| *THEMIS2* (thymocyte selection associated family member 2) | 2.29 | 0.01660 | *GREM2* (gremlin 2, DAN family BMP antagonist) | -2.52 | 0.00010 |
| *IFI44* (interferon induced protein 44) | 2.24 | 0.01944 | *TUBB3* (tubulin beta 3 class III) | -2.47 | 0.00004 |
| *ZC3H12A* (zinc finger CCCH-type containing 12A) | 2.12 | 0.02614 | *MEF2C* (myocyte enhancer factor 2C) | -2.32 | 0.03682 |
| *TNFAIP6* (TNF alpha induced protein 6) | 2.03 | 0.01012 | *CTSK* (cathepsin K) | -2.22 | 0.04135 |
|  |  |  | *NPAS1* (neuronal PAS domain protein 1) | -2.08 | 0.04135 |
|  |  |  | *TP53I11* (tumor protein p53 inducible protein 11) | -2.04 | 0.00878 |
|  |  |  | *FMN2* (formin 2) | -2.02 | 0.04135 |

**Supplementary Table S9.** Pathway and Gene family enrichment analysis of genes differentially expressed in progenitors for Cluster 2.

|  | Name | p-value | q-value<br>Bonferroni | Hit Count in<br>Query List | Hit Count in<br>Genome | Hit in Query List |
| --- | --- | --- | --- | --- | --- | --- |
| **KEGG Pathways**<br>**enriched in**<br>**genes increased**<br>**in Cluster 2**<br>**progenitors** | IL-17 signaling pathway | 1.38E-06 | 2.68E-04 | 4 | 93 | CSF3,IL1B,PTGS2,CXCL8 |
|  | Cytokine-cytokine receptor<br>interaction | 3.23E-06 | 6.27E-04 | 5 | 270 | CSF3,IL1B,IL11,LIF,CXCL8 |
|  | NOD-like receptor signaling<br>pathway | 1.53E-05 | 2.96E-03 | 4 | 170 | IL1B,OAS1,IRF7,CXCL8 |
|  | NF-kappa B signaling pathway | 9.01E-05 | 1.75E-02 | 3 | 95 | IL1B,PTGS2,CXCL8 |
| **Gene families**<br>**(genenames.org)**<br>**enriched in**<br>**genes decreased**<br>**in Cluster 2**<br>**progenitors** | Chloride channel accessory | 3.51E-03 | 6.68E-02 | 1 | 4 | CLCA2 |
|  | Myocyte enhancer factor 2<br>proteins\|MADS box family | 3.51E-03 | 6.68E-02 | 1 | 4 | MEF2C |
|  | Basic helix-loop-helix proteins | 4.11E-03 | 7.81E-02 | 2 | 110 | ID1,NPAS1 |
|  | Tudor domain<br>containing\|Spindlin family | 4.39E-03 | 8.34E-02 | 1 | 5 | SPIN2A |
|  | Actins | 5.27E-03 | 1.00E-01 | 1 | 6 | ACTC1 |
|  | DAN family | 6.14E-03 | 1.17E-01 | 1 | 7 | GREM2 |
|  | Vesicle associated membrane<br>proteins | 6.14E-03 | 1.17E-01 | 1 | 7 | VAMP8 |
|  | Small heat shock proteins | 9.63E-03 | 1.83E-01 | 1 | 11 | HSPB7 |
|  | Cathepsins | 1.31E-02 | 2.49E-01 | 1 | 15 | CTSK |
|  | Tubulins | 2.26E-02 | 4.30E-01 | 1 | 26 | TUBB3 |

**Supplementary Table S10.**
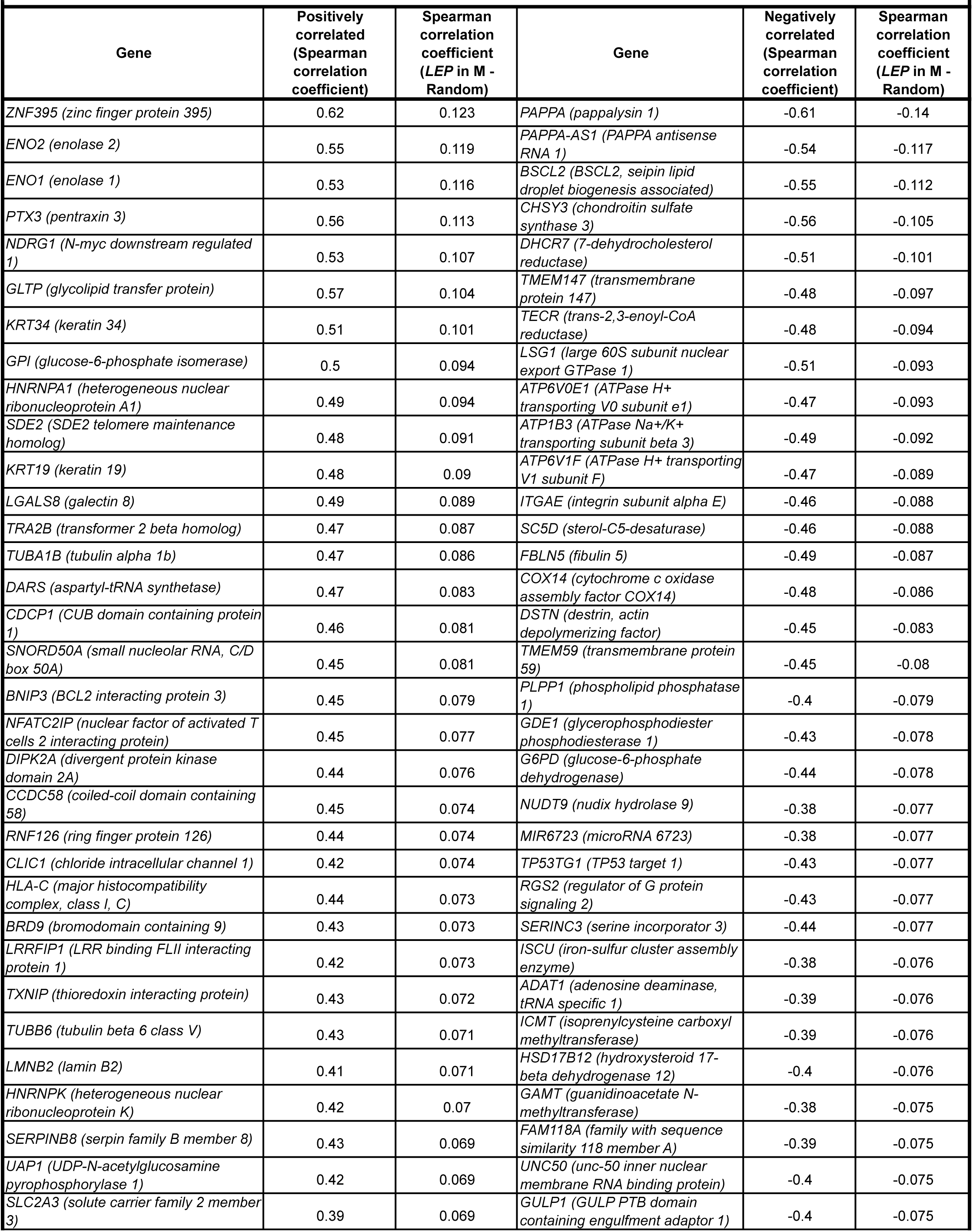
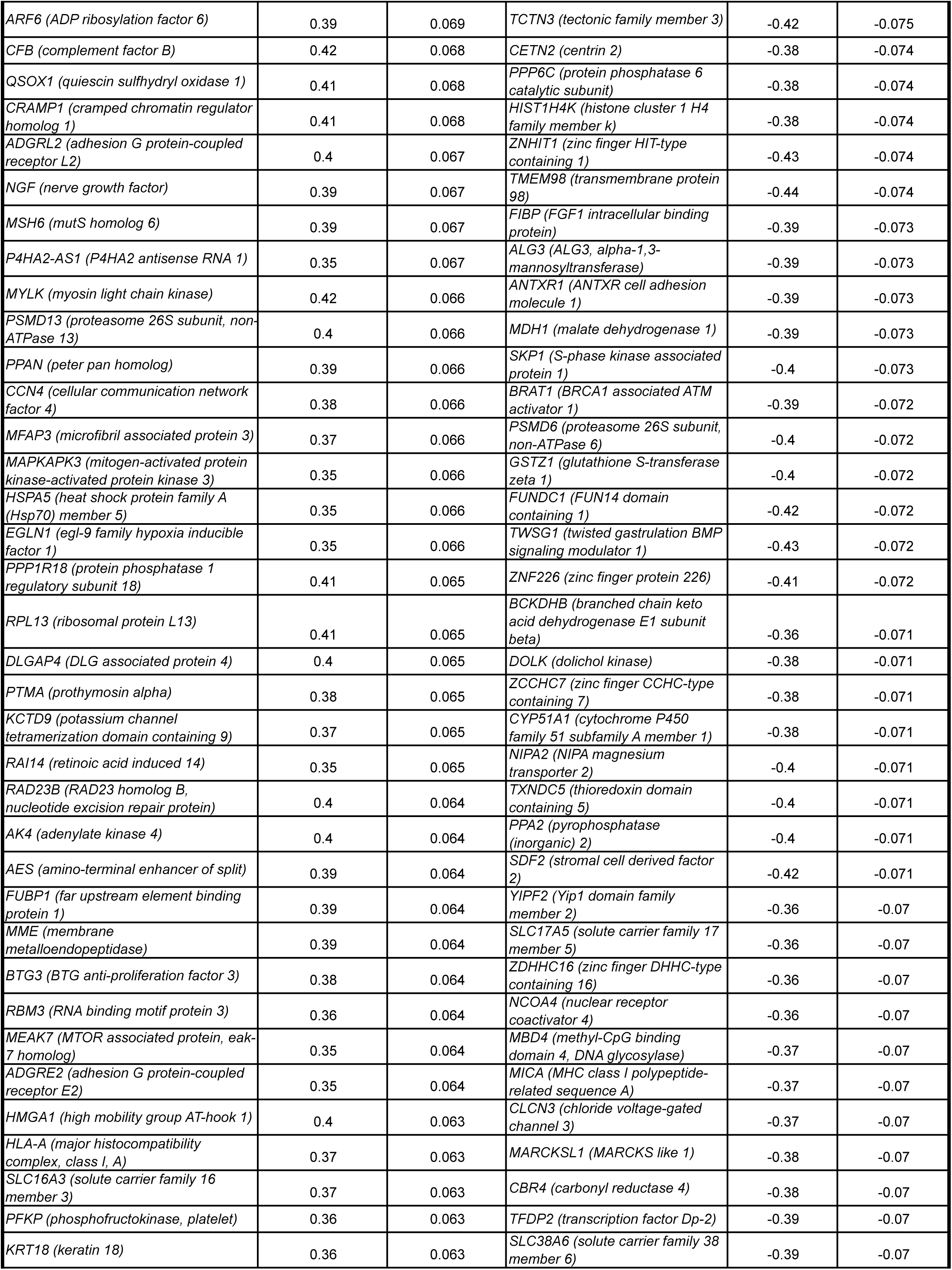
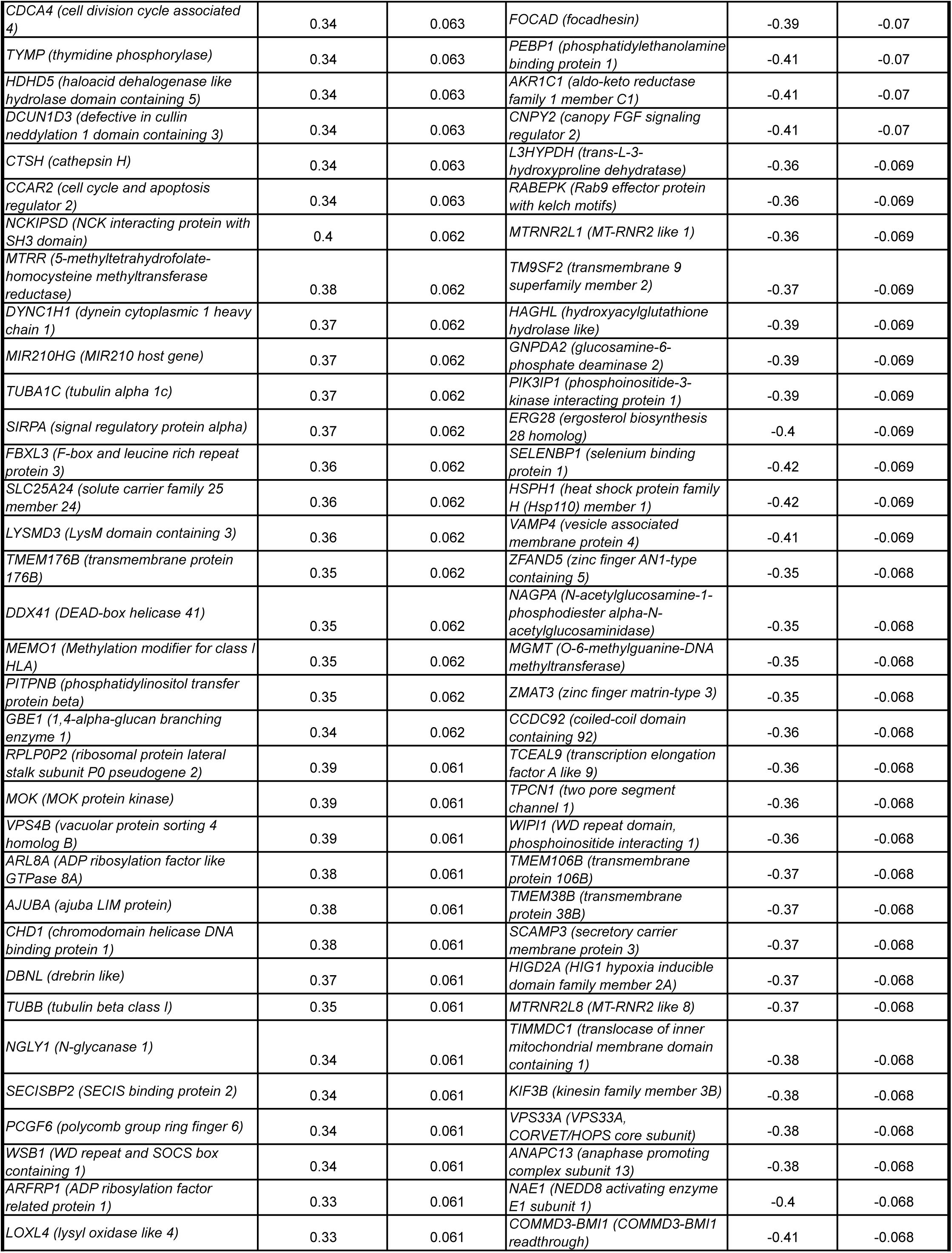
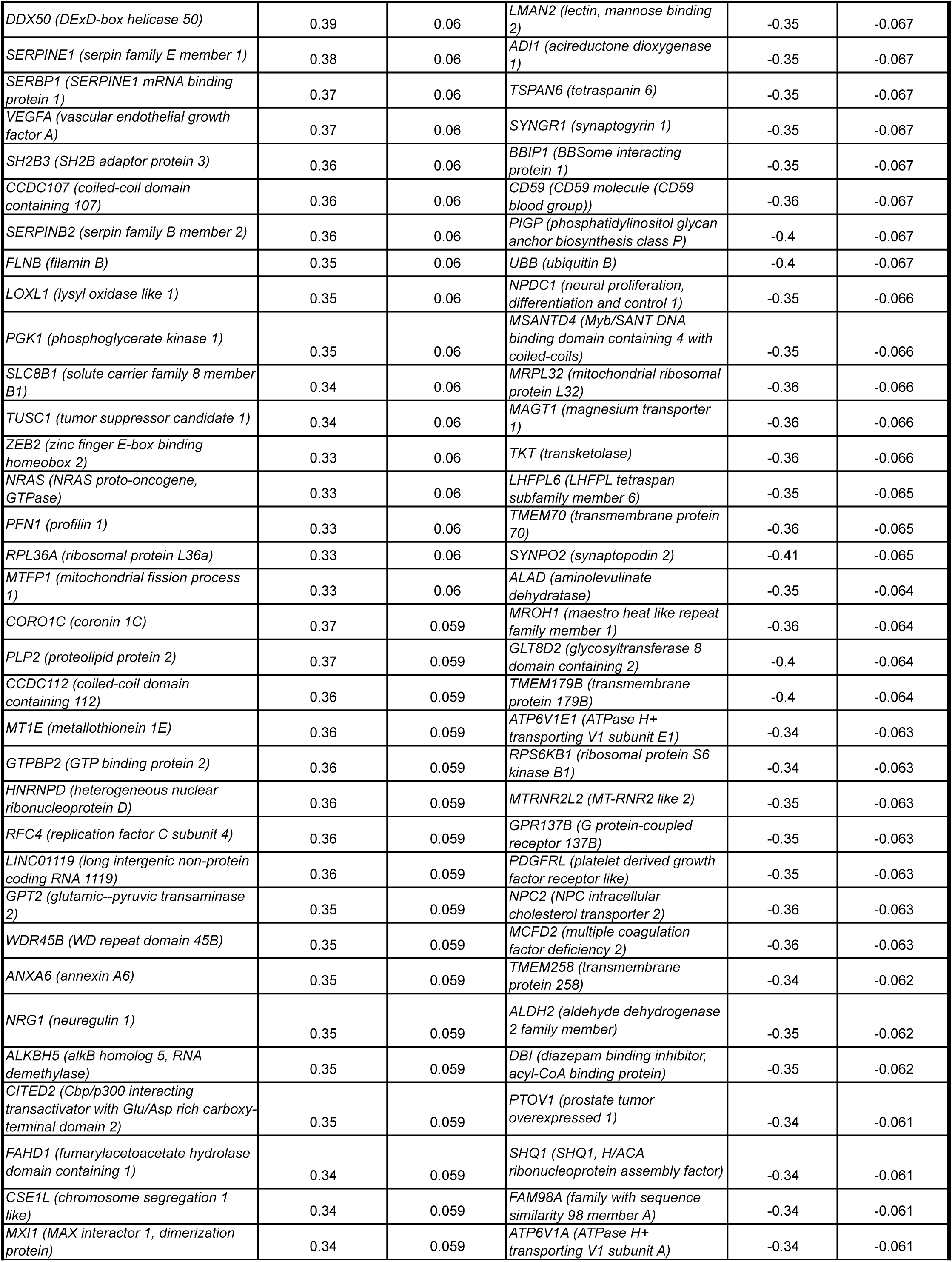
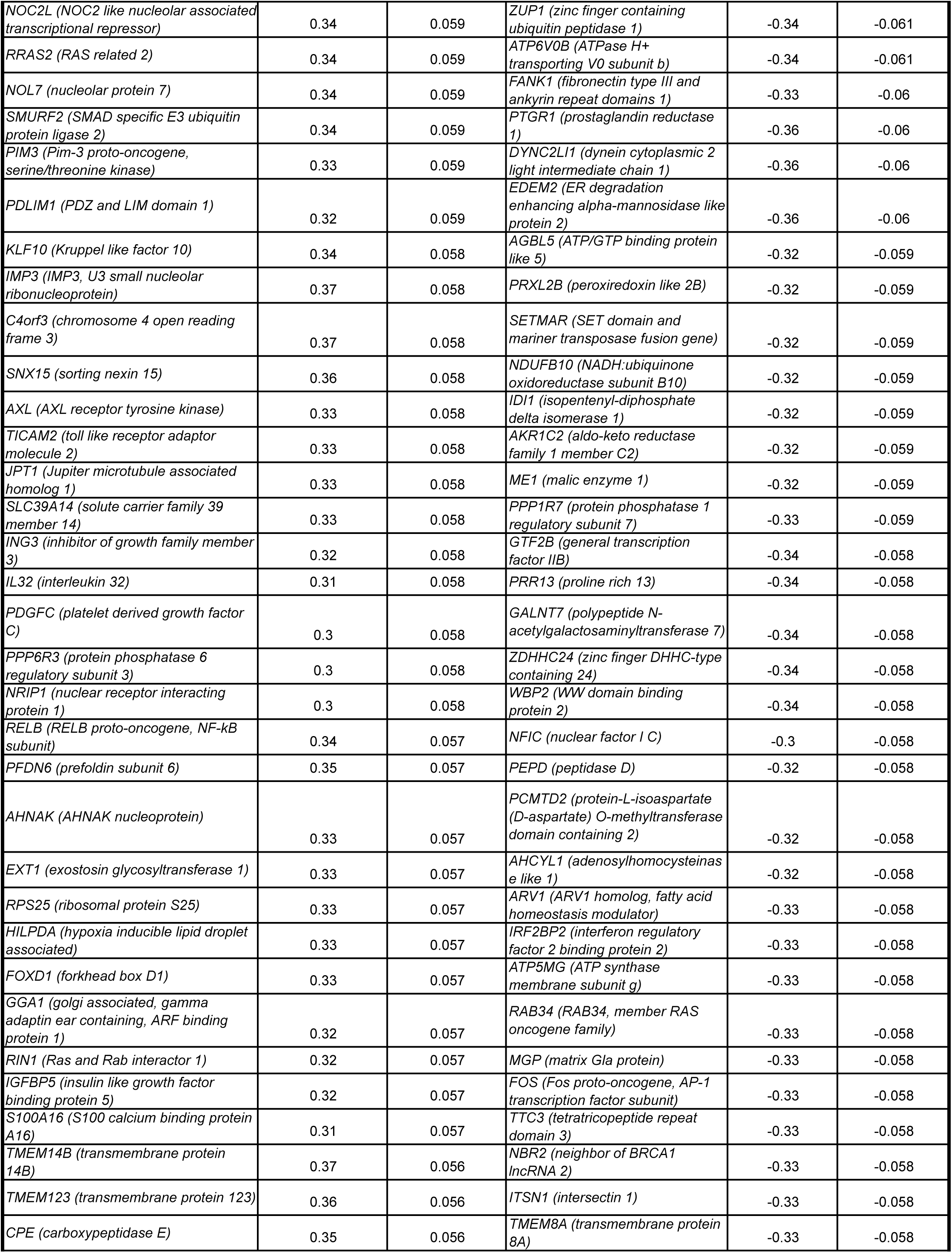
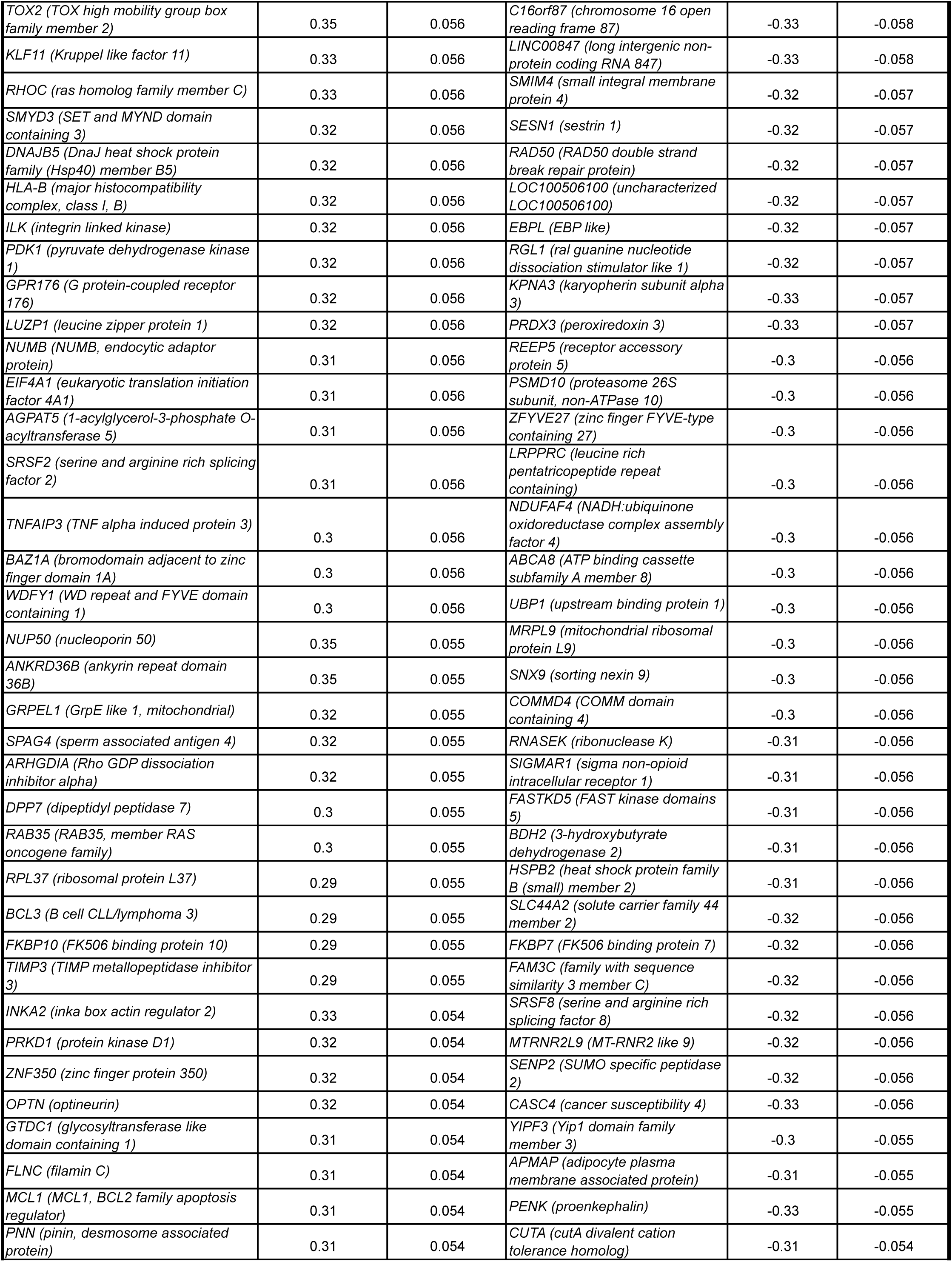
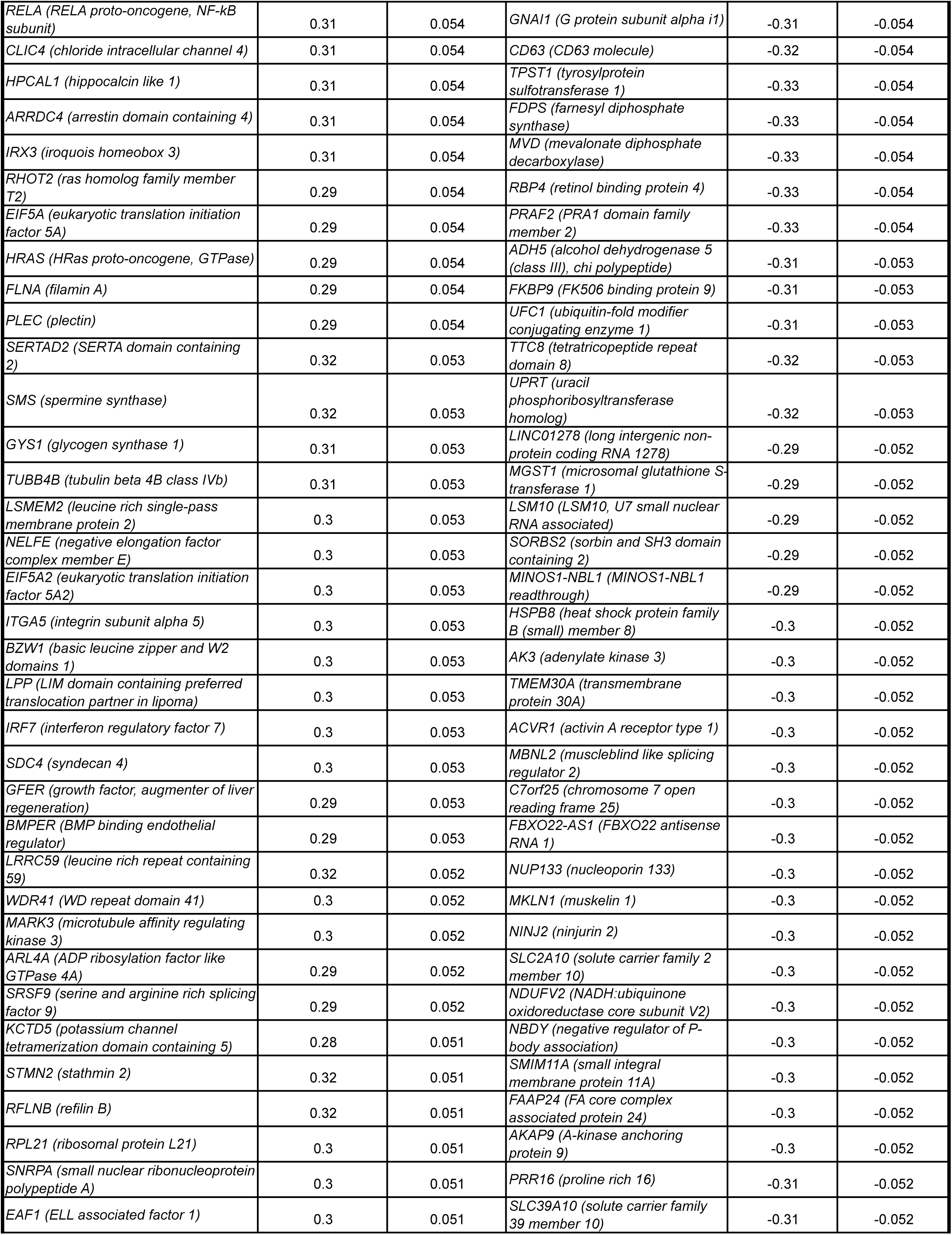
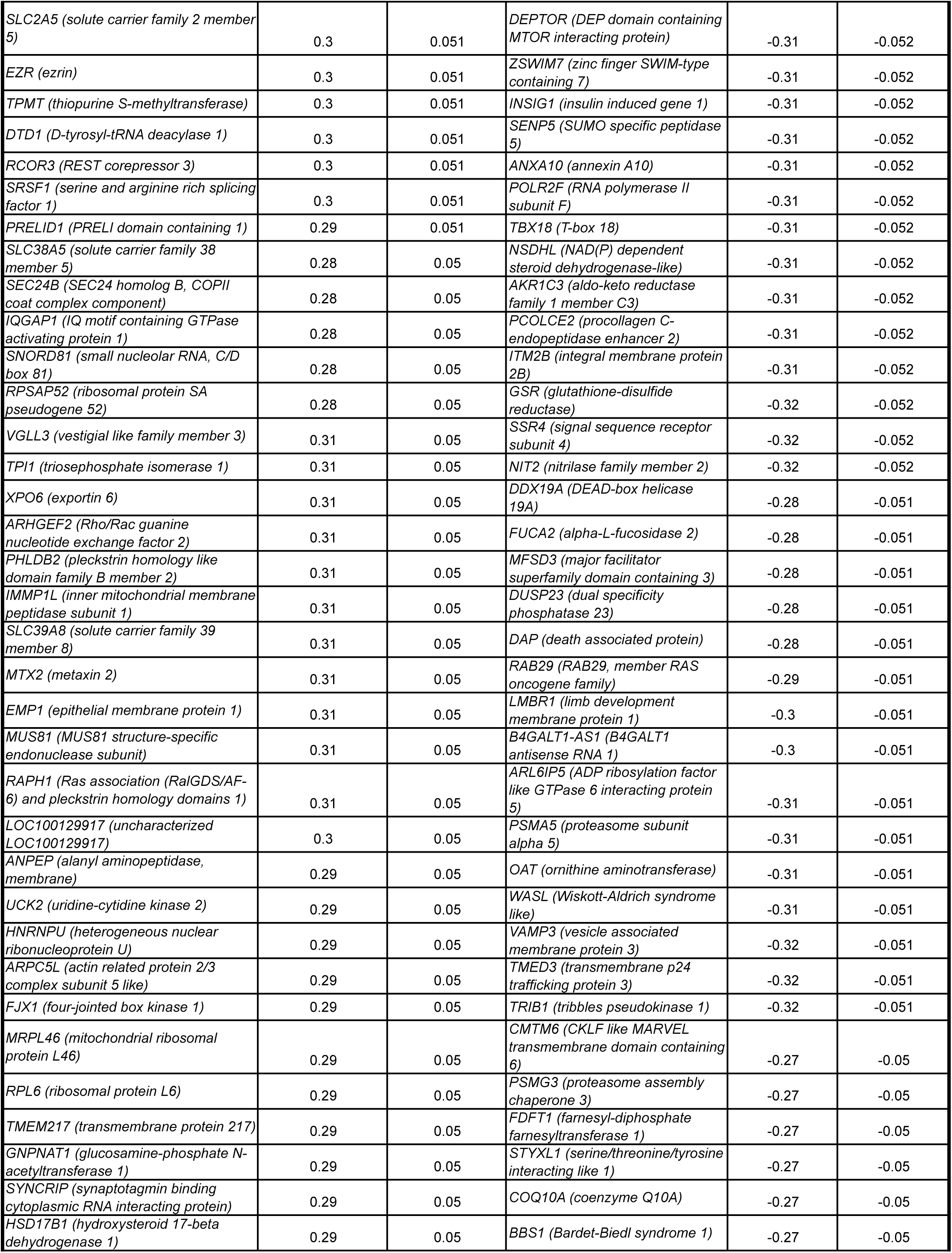
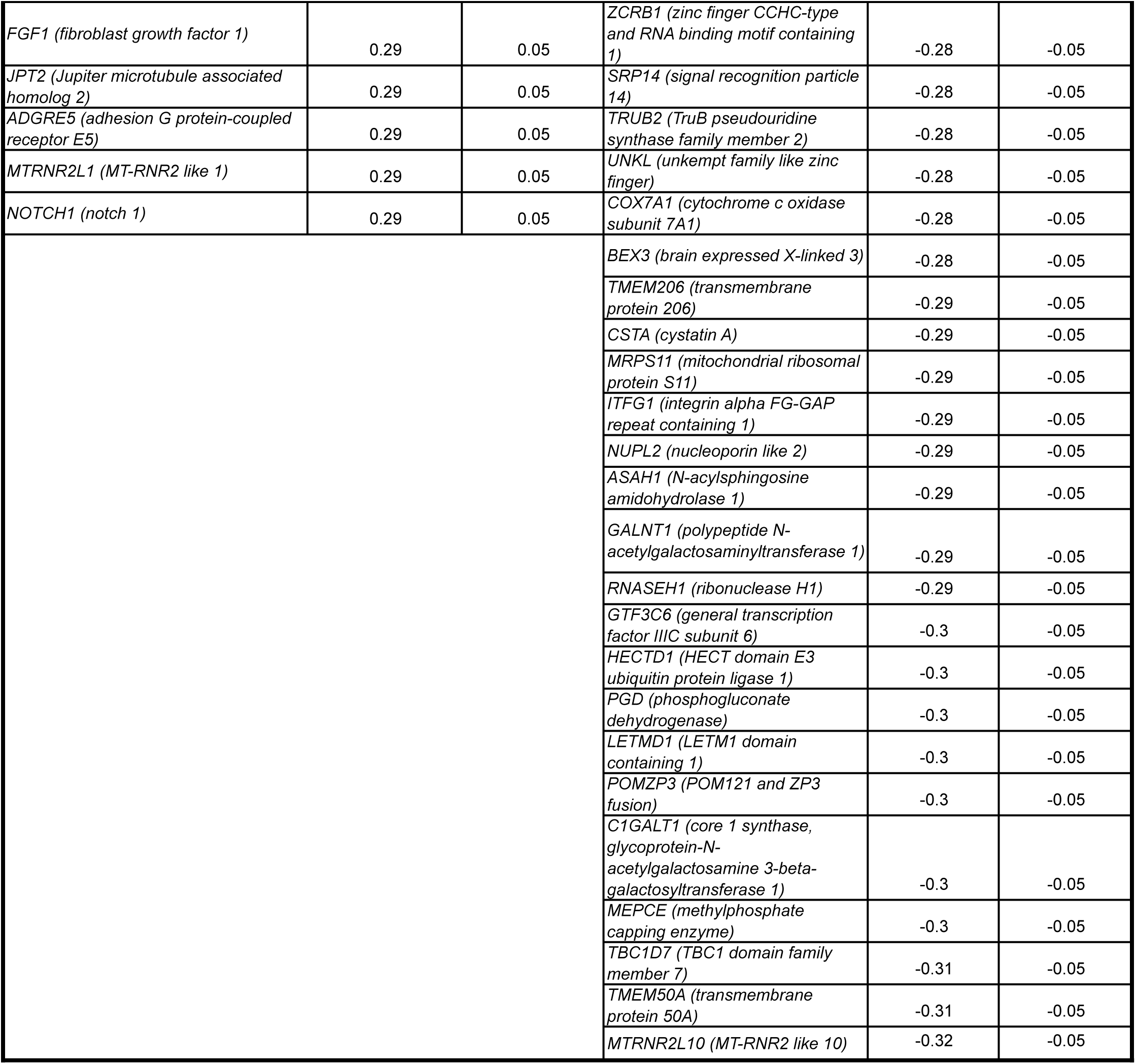
Genes in progenitors correlated with *LEP* levels in M condition

**Supplementary Table S11.**
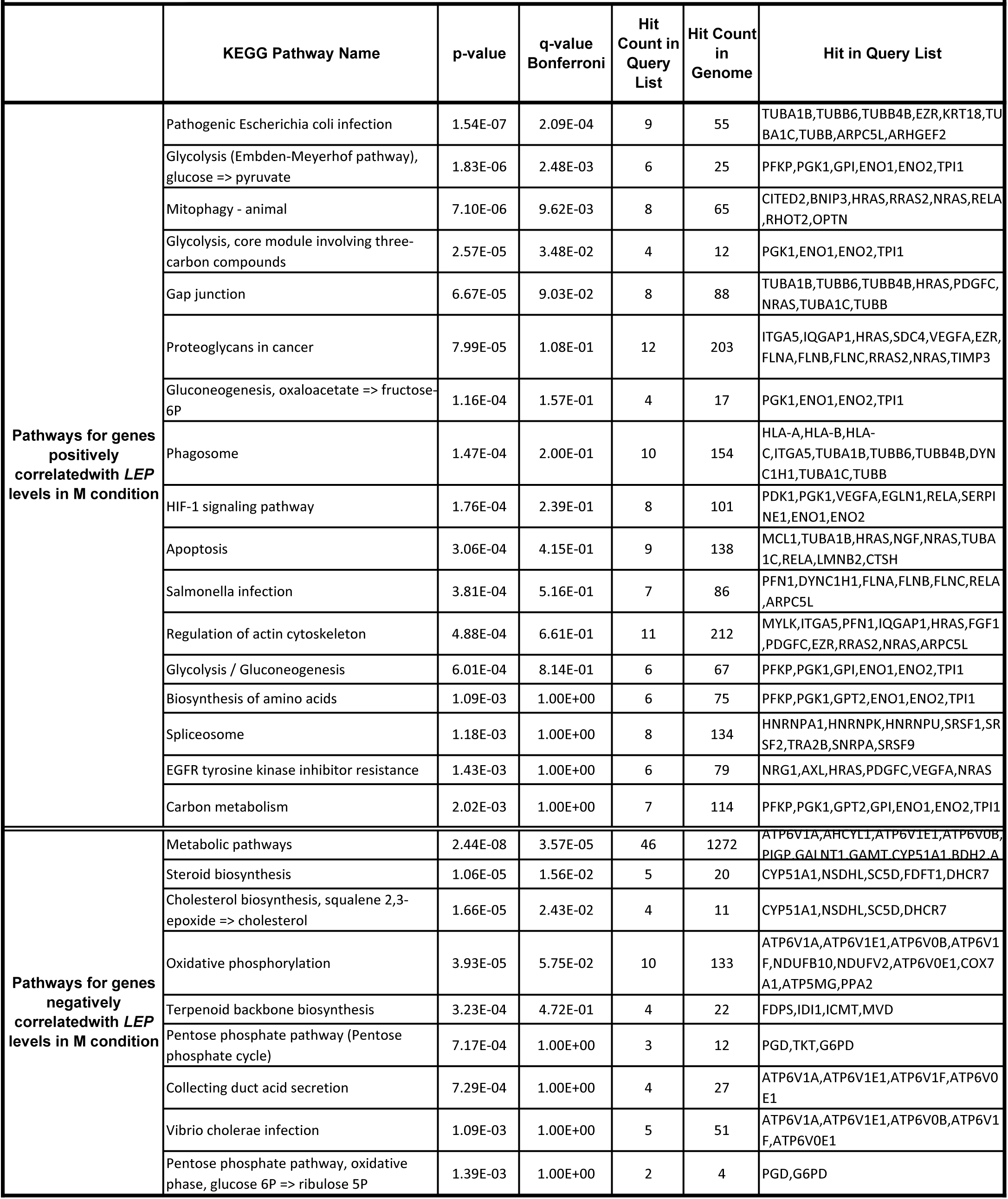
Pathway enrichment analysis of genes in progenitors correlated with *LEP* levels in M condition.

